# Semi-synthetic nanobody-ligand conjugates exhibit tunable signaling properties and enhanced transcriptional outputs at neurokinin receptor-1

**DOI:** 10.1101/2023.10.08.561411

**Authors:** Nayara Braga Emidio, Ross W. Cheloha

## Abstract

Antibodies have proven highly valuable for therapeutic development; however, they are typically poor candidates for applications that require activation of G protein-coupled receptors (GPCRs), the largest collection of targets for clinically approved drugs. Nanobodies (Nbs), the smallest antibody fragments retaining full antigen-binding capacity, have emerged as promising tools for pharmacologic applications, including GPCR modulation. Past work has shown that conjugation of Nbs with ligands can provide GPCR agonists that exhibit improved activity and selectivity compared to their parent ligands. The neurokinin-1 receptor (NK1R), a GPCR targeted for the treatment of pain, is activated by peptide agonists such as Substance P (SP) and neurokinin A (NKA), which induce signaling through multiple pathways (G_s_, G_q_ and β-arrestin). In this study, we investigated whether conjugating NK1R ligands with Nbs that bind to a separate location on the receptor would provide chimeric compounds with distinctive signaling properties. We employed sortase A-mediated ligation to generate several conjugates consisting of Nbs linked to NK1R ligands. Many of these conjugates exhibited divergent and unexpected signaling properties and transcriptional outputs. For example, some Nb-NKA conjugates showed enhanced receptor binding capacity, high potency partial agonism, prolonged cAMP production, and an increase in transcriptional output associated with G_s_ signaling; whereas other conjugates were virtually inactive. Nanobody conjugation caused only minor alterations in ligand-induced upstream G_q_ signaling with unexpected enhancements in transcriptional (downstream) responses. Our findings underscore the potential of nanobody conjugation for providing compounds with advantageous properties such as biased agonism, prolonged duration of action, and enhanced transcriptional responses. These compounds hold promise not only for facilitating fundamental research on GPCR signal transduction mechanisms but also for the development of more potent and enduring therapeutics.

## Introduction

Nanobodies (Nbs) are the smallest antibody fragments (∼ 15 kDa) that retain full capacity for binding antigens. These molecules have attracted attention as an alternative to traditional antibodies due to certain favorable characteristics, e.g., high-yield recombinant expression in bacteria, stability under conditions inhospitable to full-size antibodies, and robust methods for linkage with various cargos.^1,2^ These properties mark nanobodies as an ideal class of biomolecules for research and therapeutic applications, especially for use as modular building blocks for assembling multi-specific constructs.^2^ Nanobodies have emerged as important research tools for studying G protein-coupled receptors (GPCRs), the largest and most diverse group of membrane receptors in humans and target of more than 30% of the drugs on the market.^2-5^ Receptor-specific nanobodies are widely used to detect or enforce specific GPCR conformational states for mechanistic and structural studies, respectively.^5-7^ Nanobodies have also been exploited as diagnostics and therapeutic tools.^2,4^

The linkage of peptide agonists of GPCRs with Nbs can yield conjugates with improved signaling potency and selectivity compared to free peptides.^8^ For example, the conjugation of weakly active fragments of parathyroid hormone (PTH) to nanobodies that bound the type-1 PTH-receptor (PTHR1), or epitope tags incorporated therein, provided conjugates with significantly improved biological potencies (up to 7800-fold).^8,9^ These conjugates also showed improved specificity for a single receptor subtype relative to the index ligand from which they were derived.

A separate metric of ligand performance relates to the types of signaling responses induced upon ligand application. Ligands often exhibit signaling through more than one receptor-coupled pathway and individual ligands induce a characteristic balance of signal flux through each of these pathways. Past work has shown that variation in ligand structure can alter the profile of signaling responses induced. Here we sought to evaluate whether the linkage of ligands to Nbs would generate conjugates that show distinct signaling outputs compared to natural ligands. To test this, we used ligands of the neurokinin-1 receptor (NK1R).

The NK1R is a GPCR from the tachykinin receptor family that is targeted by peptide agonists. Substance P (SP), an 11–mer peptide amidated at its *C*-terminus (RPKPQQFFGLM), is the most selective NK1R agonist.^10-12^ However, other endogenous agonists, such as neurokinin A (NKA; HKTDSFVGLM) also bind NK1R, albeit with lower affinity compared to SP.^10-12^ GPCRs including NK1R receive various extracellular signals and convert them into cellular responses by activating associated G-proteins, β-arrestins, and other downstream effectors.^10,12,13^

We provide evidence of the capacity of Nb-peptide conjugates to induce NK1R-mediated signaling and transcriptional response profiles that differ from, and are sometimes superior to, the prototype peptide ligand. By exploring the unique properties of Nb-peptide conjugates, we aim to advance our understanding of GPCR signaling and facilitate the development of therapeutic candidates with useful properties.

## Results

### Receptor constructs and synthesis of Nb-peptide conjugates

Since at current there are no nanobodies that directly bind NK1R, we inserted a sequence comprised of three short epitope tags **(Table S1)** into the extracellular N-terminus of NK1R to enable recognition by the nanobodies Nb_6e_, Nb_alfa_, Nb_BC2_ .^14-17^ We established stable cell lines expressing epitope-tagged NK1R **(Figure 1A)**. To target NK1R, we developed conjugates composed of Nbs and NKA or a truncated version of SP (SP_6-11_). In consideration of previous studies with the PTHR1 system^8^, we opted for ligands with modest binding, instead of the tightest binding peptide agonist (substance P), as weaker binding ligands often exhibit more significant changes in properties upon Nb linkage. We expressed the nanobodies Nb_6e_, Nb_alfa_, Nb_BC2_ and the negative control Nb_GFP_ in *Escherichia coli* (*E. coli*) with a hexahistidine tag (6× His-tag) and a sortase A recognition motif (LPETG) at the *C*-terminus. The nanobodies were purified using nickel-NTA sepharose beads followed by size exclusion chromatography. We synthesized NKA and SP_6-11_ along with their analogues with an *N*-terminal triglycine extension (G3NKA, G3SP_6-11_) by conventional Fmoc-SPPS (9-fluorenymethyloxycarbonyl-solid phase peptide synthesis). The peptides were purified by reversed-phase (RP)-HPLC and characterized by mass-spectrometry (MS) **(Figure S1)**. The addition of a triglycine motif to the *N*-terminus of these tachykinins was not expected to substantially alter their activity, as the region responsible for receptor activation resides in the *C*-terminus (FXGLM-NH2), while the N-terminal region contributes to receptor subtype selectivity.^18^ The activities of NKA and SP_6-11_ were compared to their triglycine extended analogues. This comparison showed modest effects for triglycine extension, except for G3SP_6-11_ (EC_50_ 0.3 nM), which was almost 100 times more potent than SP_6-11_ (27.5 nM) in signaling through the G_q_ pathway **(Figure S2)**. We employed sortase A-mediated labeling (sortagging) to conjugate G3NKA or G3SP_6-11_ to the nanobodies (**Figure 1B**), followed by purification and analysis by MS **(Figure S3 & Table S2)**.^19, 20^

**Figure 1.**
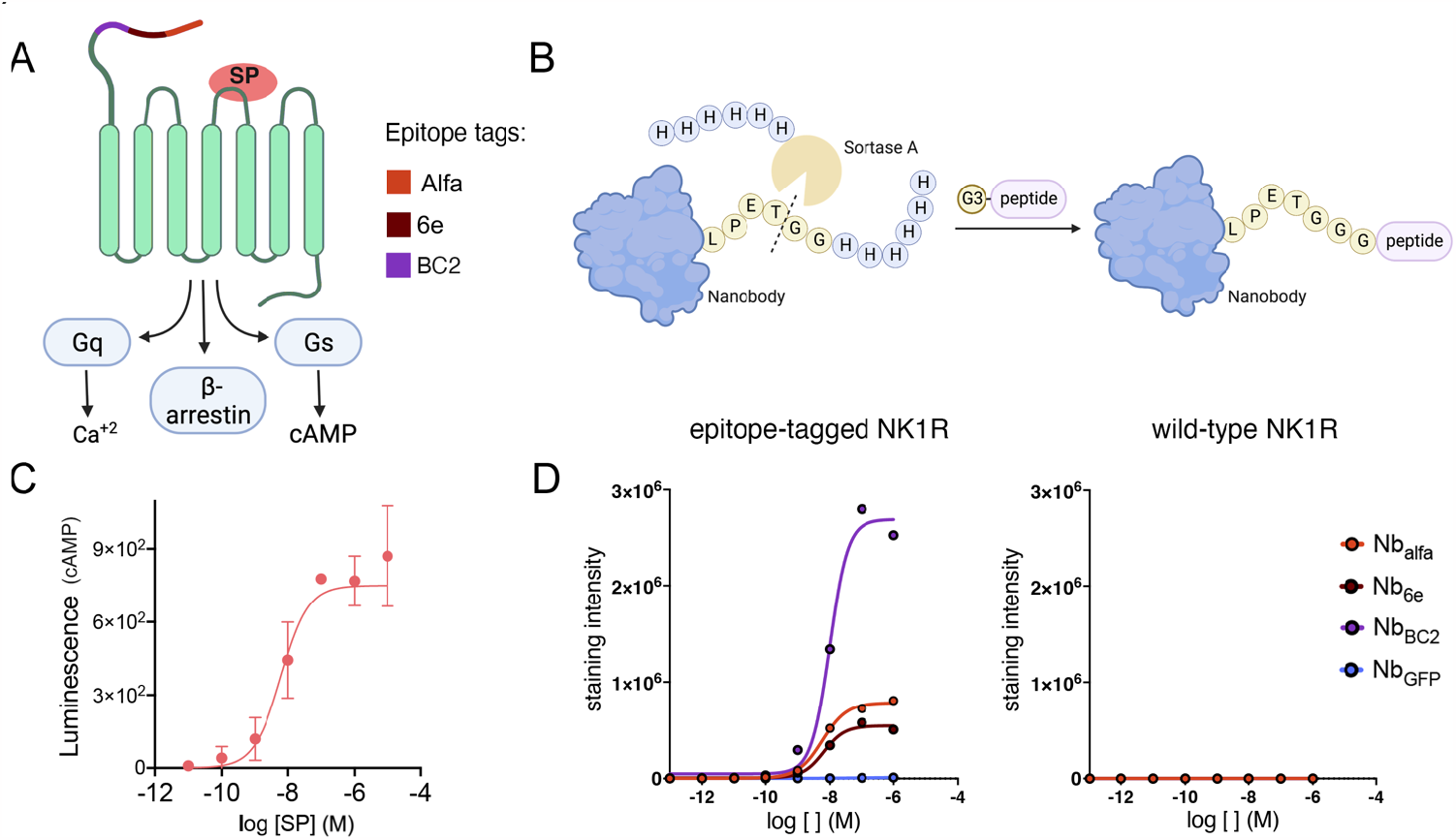
Receptor constructs and synthesis of Nb-peptide conjugates. **(A)** NK1R signals through G_q_, G_s_ (cAMP production) and β-arrestin. Epitope tags (alfa, 6e and BC2) were inserted into the *N*-terminus to enable Nb recognition. (**B)** Nanobodies and peptides were conjugated via Sortase A (SrtA)-mediated ligation. (**C)** Substance P (SP) activation of epitope-tagged NK1R. Receptor activation was evaluated by signal generated from a stably expressed cAMP-responsive luciferase construct. Data points (mean ± SD) correspond to technical replicates from a single representative experiment. Data from independent replicates are shown in **Figure S4**. (**D)** Biotin-labeled nanobodies, except Nb_GFP_, bound to epitope-tagged NK1R, while no binding was observed with NK1R wild-type. Dose-response curves were generated from application of a sigmoidal concentration-response 3 parameter model. Data from independent replicates are shown in **Figure S5**.

We assessed the functionality of epitope-tagged NK1R by evaluating the capacity of SP to induce the production of cyclic adenosine monophosphate (cAMP), a second messenger molecule generated upon NK1R activation and signaling through G_s_. We used a cell line that stably expresses epitope-tagged NK1R and a luciferase-based cAMP-responsive reporter.^21^ SP exhibited a concentration-dependent induction of cAMP production (EC_50_ 17 ± 3 nM), confirming the functionality of the engineered receptor (**Figure 1C & S4**). Additionally, we used flow cytometry to assess the binding of Nbs to tagged receptors on live cells. As a control, we also included Nb_GFP_ which targets GFP and, therefore, is not expected to bind to cells expressing epitope-tagged NK1R. Nb_6e_, Nb_alfa_, Nb_BC2_ bound to the cell lines expressing epitope-tagged NK1R while Nb_GFP_ did not (**Figure 1D**). Nb_6e_ and Nb_alfa_ displayed similar staining intensity and potency, while Nb_BC2_ exhibited more intense staining with similar potency (**Figure 1D & S5**). The origin of the difference in staining intensity between the different tag-specific nanobodies is unknown but it might be related to the proximity of the tag to the receptor transmembrane domain. As expected, none of the nanobodies bound to cells expressing NK1R lacking epitope tags (wild-type; NK1R WT) (**Figure 1D**).

### Conjugation of NK1R ligands with nanobodies imparts specialized signaling properties

Next, we sought to comprehensively evaluate the pharmacological properties of the conjugates. Activation of NK1R by NKA and SP_6-11_ induces intracellular signaling cascades mediated by G_q_, G_s_ and β-arrestin (**Figure 1A**).^10, 22, 23^ First, we assessed the ability of the conjugates to trigger the G_s_ pathway by measuring cAMP production using a luciferase-complementation (Glosensor) assay. Nb-NKA and Nb-SP_6-11_ conjugates exhibited modest to weak efficacy for stimulating cAMP production on cells expressing epitope-tagged NK1R (**Figure 2A**). Notably, conjugate agonist behavior was dependent on the identity of the nanobody linked with peptide ligand. Nb_alfa_- and Nb_6E_-NKA conjugates exhibited high potency, low to modest efficacy activation of cAMP signaling; Nb_BC2_-NKA exhibited low efficacy; and Nb_GFP_-NKA conjugate was essentially inactive (**Figure 2A**). This pattern of agonism of G_s_ signaling for conjugates does not correlate with the binding performance of tag-binding (or control) nanobodies used to prepare these conjugates **(Figure 1D)**. The activity of Nb-NKA conjugates was uniformly weak in G_s_ signaling on cell lines expressing NK1R WT, indicating that the linkage of ligands to nanobodies may hinder the necessary ligand-receptor interaction needed for subsequent G_s_ coupling when the nanobody epitopes are absent from the receptor of interest (**Figure S6**). Nb-SP_6-11_ conjugates failed to elicit cAMP production on both WT and epitope-tagged NK1R cell lines (**Figure 2B & S6)**.

**Figure 2.**
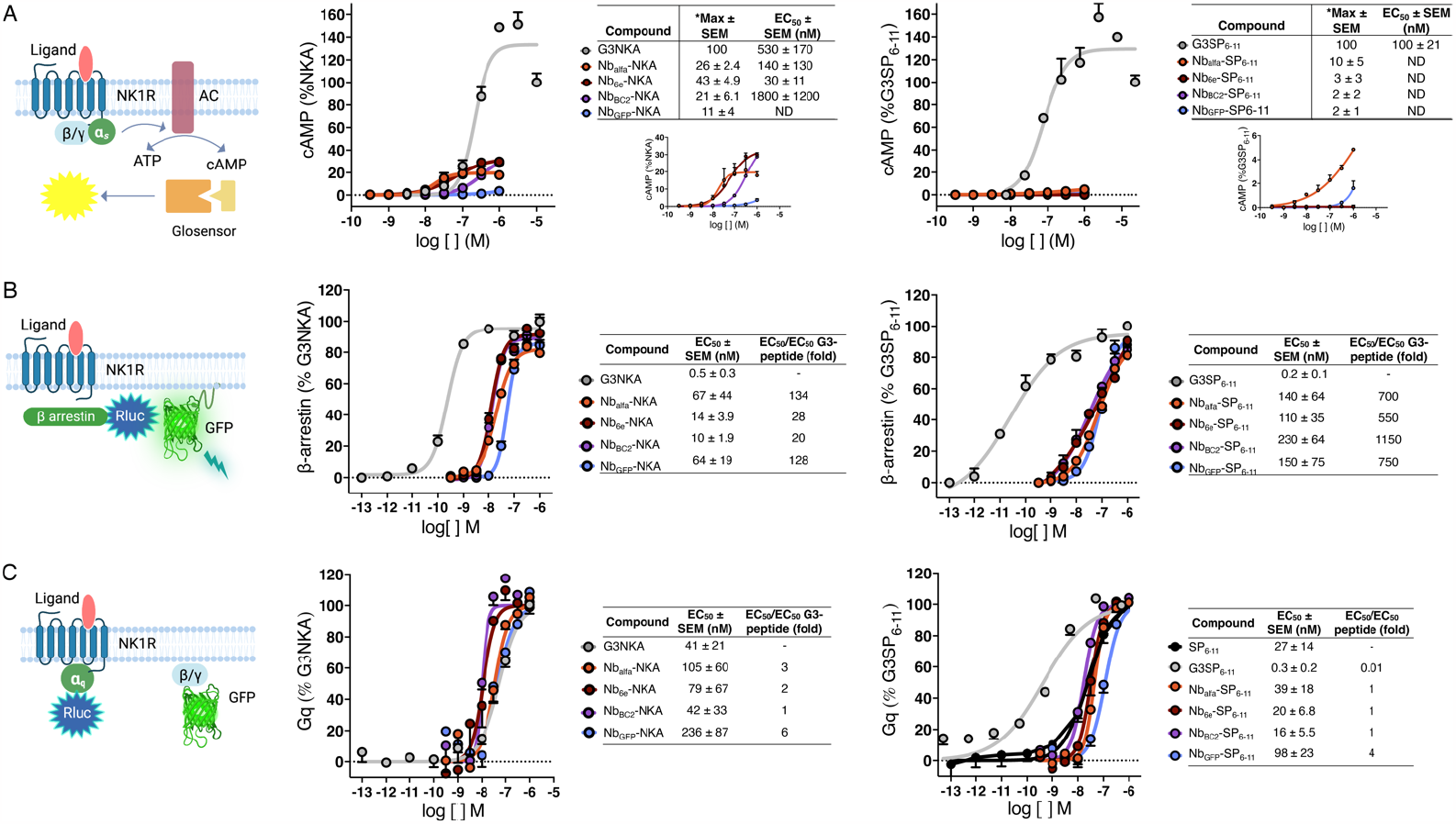
Effects of peptides and conjugates on NK1R signaling. **(A)** HEK293 cells stably transfected with Glosensor cAMP reporter (Promega Corp.)^21^ and epitope-tagged were treated with varying concentrations (1 pM – 10 μM) of the indicated peptides or conjugates. Activation was assessed by cAMP production after 6 min as described in Methods. Insets of concentration-responses without G3-peptides included are shown (under table) to allow better visualization of the activity of the weaker compounds. *Max activity values were calculated by normalizing the response at 1 μM for conjugates to that of G3NKA or G3SP_6-11_ at 10 μM (n=3). EC_50_ for conjugates with maximal responses lower than 30% were not determined (ND). **(B)** HEK293 cells stably expressing epitope-tagged NK1R were transfected with Gq TRUPATH Gα/β/γ biosensor plasmid and treated with varying concentrations (0.1 pM – 1 μM) of the indicated peptides or conjugates and BRET signals were measured. **(C)** HEK293 cells stably expressing epitope-tagged NK1R were transfected with β-arrestin2-RlucII/rGFP-CAAX plasmids and treated with varying concentrations (0.1 pM – 1 μM). Representative concentration-response curves are shown, where data points correspond to mean ± SD from technical replicates. EC_50_ values were calculated from the fitting of a sigmoidal concentration-response model to data from n ≥ 3 independent experiments.

Then, we performed BRET assays using a G_q_ TRUPATH Gα/β/γ biosensor plasmid^24^, which reports on the separation of G_α_ subunit from G_βγ_ heterodimer upon ligand-induced activation. Using this approach, we evaluated G_q_ activation using the clonal cell line expressing epitope tagged NK1R used for all other assays. Nb-NKA and Nb-SP_6-11_ conjugates exhibited full efficacy G_q_ pathway signaling with modest reductions in signaling potency relative to comparator peptides **(Figure 2B)**. This reduction was less pronounced for Nb-NKA conjugates (< 6-fold difference in EC_50_ values) than those seen with Nb-SP_6-11_ conjugates (> 60-fold difference) **(Figure 2B)**. There was little difference between conjugates comprised of Nbs that bound to the tagged receptor versus negative control Nbs. We also made the surprising observation that the presence of the three *N*-terminal glycines in G3SP_6-11_ was responsible for an increase in potency of almost 100-fold relative to SP_6-11_ alone in this assay **(Figure S2)**. Thus, Nb-G3SP_6-11_ conjugates performed similarly to SP_6-11_ (but not G3SP_6-11_) in signaling through the G_q_ pathway (< 4-fold difference in EC_50_). In line with these findings, we observed that conjugates and index peptides were similarly efficacious for inducing intracellular calcium mobilization, a downstream response mediated by G_q_ activation (**Figure S7**). These results collectively demonstrate that the Nb conjugation and the Nb-epitope interactions play a minor role in modulating the NKA/SP_6-11_-NK1R interaction that triggers G_q_ signaling.

The function of NK1R is mediated in part by the ligand-induced recruitment of β-arrestins (βarrs), intracellular proteins that prevent further receptor–G protein coupling, recruit phosphodiesterases to the cell surface, and promote receptor internalization.^13^ We evaluated the ability of Nb-NKA and Nb-SP_6-11_ conjugates to induce the recruitment of β-arrestin2 in a bioluminescence resonance energy transfer (BRET) assay.^25^ These assays were performed using the same clonal HEK293 cell lines used for cAMP induction and G_q_ activation assays, enabling consistency in receptor expression levels across experiments. Nb-NKA conjugates displayed the same efficacy as G3NKA, although reductions (20-130-fold) in potency were observed (**Figure 2C**). The negative control Nb_GFP_-NKA conjugate was comparable in potency to other Nb-NKA conjugates for stimulating recruitment of βarr2, suggesting a minimal role for nanobody-tag interactions in modulating βarr2 recruitment. Nanobody conjugation had an even greater impact on SP_6-11_ behavior, reducing potency by at least 500-fold relative to the index peptide **(Figure 2C)**.

Collectively, these data show that the activity pattern of the conjugates varies depending on the pathway, ligand, and nanobody identity **(Figure 2)**. One striking observation is that Nb-SP_6-11_ conjugates exhibited full efficacy for stimulation of G_q_ and βarr recruitment, which contrasts their inactivity in cAMP induction assays. These observations indicate a complex connection between nanobody binding, receptor activation triggered by the conjugates, and intracellular signaling.

### Nb conjugation potentiates ligand binding at NK1R orthosteric site and promotes prolonged cAMP production

We evaluated the binding of Nb-peptide conjugates to epitope-tagged NK1R using a competition assay analyzed by flow cytometry. Cells expressing epitope-tagged NK1R were incubated with 30 nM of fluorescently labeled SP (SP-AF647; **Figure S8A**) and varied concentrations of unlabeled conjugates or free peptides. Nb-NKA conjugates (100 nM) comprised of Nbs that target the receptor, especially Nb_6e_ and Nb_BC2_, demonstrated enhanced performance compared to NKA alone for outcompeting SP-AF647 for binding to the target receptor (**Figure 3A & S7)**. Conjugation of SP_6-11_ to Nb_6e_, and more markedly Nb_BC2_, also augmented competitive binding activity at 100 nM concentration (**Figure 3B and S7)**. Conversely, conjugation to Nb_alfa_ and Nb_GFP_ (100 nM) significantly diminished competitive binding affinity (**Figure 3B and S8**). We hypothesize that the smaller size of SP_6-11_ renders this peptide more susceptible to conjugation-induced variation in binding. More broadly, findings indicate that nanobody-tag interactions might lead to improved association with the orthosteric site; however, caveats apply. In addition, it suggests that the proximity of the tag to the orthosteric binding site might impact binding in complex ways. For example, Nb_alfa,_ conjugates that bind to a tag located closer to the N-terminus, had a worse ability to outcompete SP than Nb_6e_ and Nb_BC2_ conjugates, despite similar performance in direct Nb binding assays **(Figure 1D)**.

**Figure 3.**
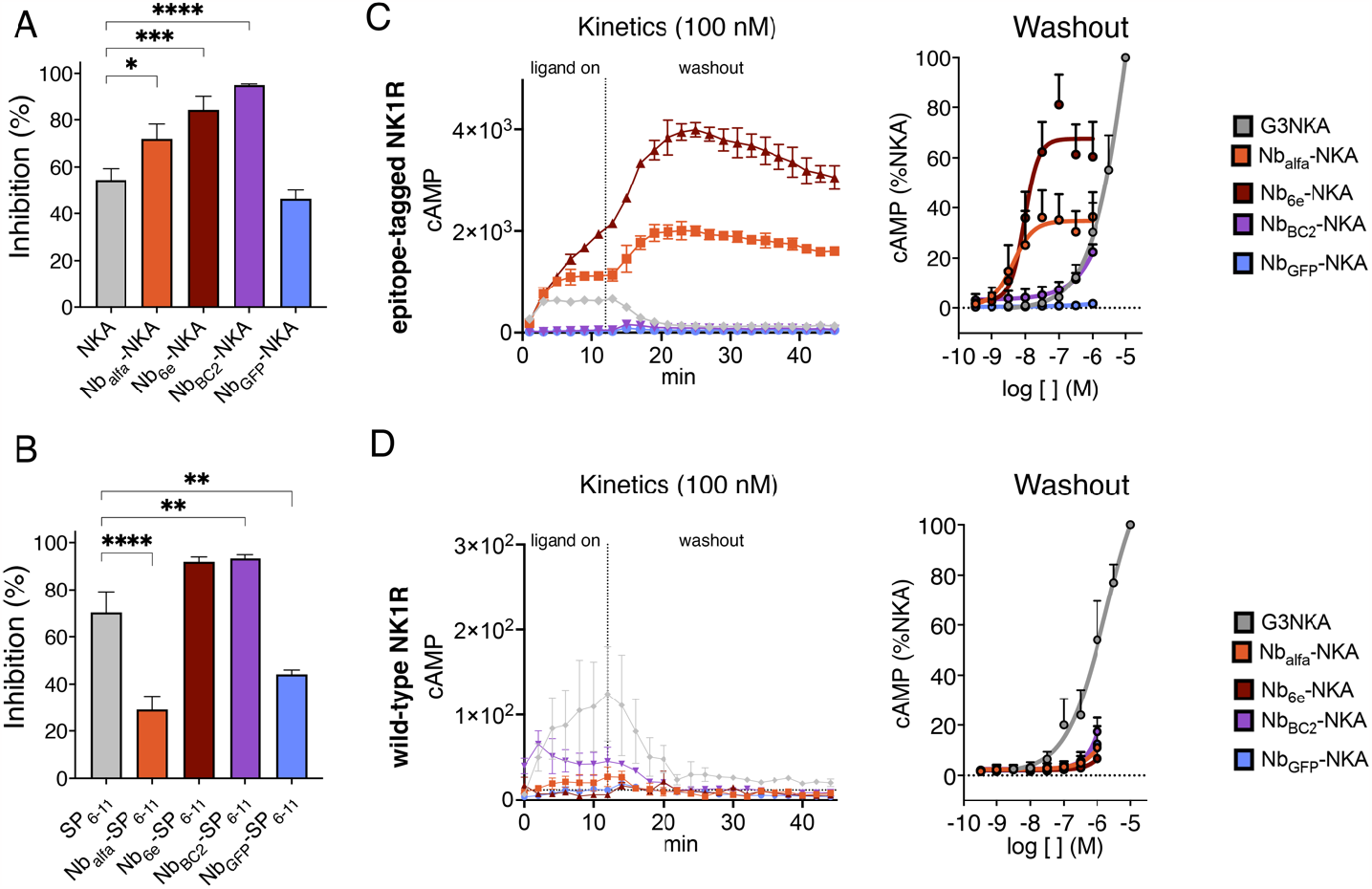
Impact of Nb conjugation on ligand competition binding and duration of cAMP activity. **(A, B)** A competition binding assay was performed using flow cytometry to assess the ability of 100 nM **(A)** NKA and Nb-NKA or **(B)** SP_6-11_ and Nb-SP_6-11_ to outcompete the binding of fluorescently labeled SP to epitope-tagged NK1R. Data represent mean ± SEM from n ≥ 3 independent experiments. (**C, D)** Representative concentration-response curves or cAMP production kinetics (mean ± SD) for washout assays. 12 minutes after ligand addition, excess/unbound ligand was removed, fresh media was added to cells expressing **(C)** epitope-tagged or **(D)** wild-type NK1R, and cAMP responses were measured for an additional 30 min (washout) (n ≥ 3) according to Methods. Washout assay data were quantified as the area under the curve (AUC). Curves result from the fitting of a sigmoidal concentration-response model to data. Differences were evaluated for statistical significance with a one-way ANOVA followed by Dunnett correction. *p < 0.03, **p < 0.002, ***p < 0.0002, **** p < 0.0001

We investigated the impact of conjugating NKA to Nbs on the duration of cAMP production. Despite reduced efficacy, Nb_alfa_-NKA and Nb_6e_-NKA stimulated more enduring activation of tagged NK1R upon removal of excess/unbound ligand (“washout”) relative to simple peptide agonists alone (**Figure 3C & S9**). In contrast, we observed a rapid washout of all Nb-NKA conjugates in NK1R WT cells (**Figure 3D**). We further hypothesized that the two-site binding mechanism of conjugates (nanobody-tag site and ligand-receptor orthosteric site) would reduce the impact of NK1R competitive antagonists that only block the receptor orthosteric site. To test this, we evaluated the durability of Nb_6e_-NKA signaling by adding the NK1R competitive antagonist spantide I^26^ during the washout. The addition of spantide I (1 μM) accelerated the dissipation of G3NKA signaling but had a smaller effect on Nb_6e_-NKA **(Figure S10)**. This finding suggests that receptor-directed nanobody tethering is a promising method for promoting enduring signaling responses even in the presence of competitive antagonists.

### Nb-NKA conjugation enhances G_s_ and G_q_-associated transcription output

Transcriptional modulation is a downstream event in GPCR signaling cascades, occurring after the activation of various intracellular signaling pathways. We hypothesized that prolonging the activation of the cAMP signaling pathway could result in amplification of downstream transcriptional responses mediated by G_s_ or other pathways. To test this, we used a transcriptional reporter assay in cells expressing tagged NK1R to measure the performance of conjugates in driving G_s_-mediated transcription. All Nb-NKA conjugates at a concentration of 35 nM, except Nb_GFP_-NKA (negative control), increased the level of G_s_-mediated transcription by at least 5-fold compared to NKA (**Figure 4**). These observations suggest that in the context of Nb-ligand conjugates, the interaction between the nanobody and the receptor can play a complex role in promoting receptor activation, sustained signaling and increased transcription for Nb-ligand conjugates.

**Figure 4.**
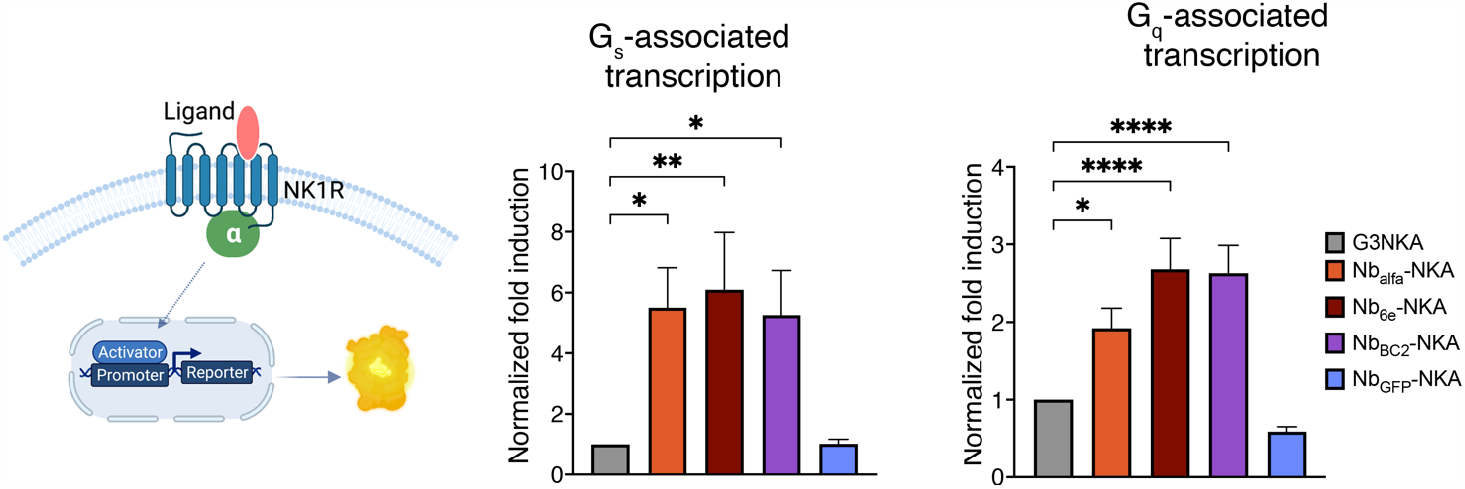
Evaluation of peptide and conjugate performance on transcription. Transcriptional responses were evaluated through transfection of cells expressing epitope tagged NK1R with a luciferase reporter plasmid reporting on G_s_ or G_q_ signaling. Cells were incubated with ∼35 nM of the indicated peptides or conjugates for 17 hours and transcription was measured as described in Methods. Data represent mean ± SEM from n ≥ 3 independent experiments. One-way ANOVA followed by Dunnett correction was performed to assess differences between free peptides and conjugates. *p < 0.03, **** p < 0.0001.

We also investigated whether Nb-tag interactions impact G_q_-mediated transcription. At a dose of 35 nM, all Nb-NKA conjugates, except Nb_GFP_-NKA (negative control), increased by at least 2-fold the magnitude of the G_q_-mediated transcriptional output compared to G3NKA (**Figure 4**). These data present trends that are clearly distinct from those observed in assays that measure upstream signaling **(Figure 1C)**. We observed negligible differences in G_q_-mediated transcriptional response induced by Nb-SP_6-11_ compared to SP_6-11_ **(Figure S11)**.

## Discussion

The Nb-ligand tools generated in this study have facilitated the investigation of three distinct lines of inquiry concerning NK1R signaling and the design of GPCR ligands in a broader context. First, we explored the consequences of fusing an NK1R ligand to a larger protein (Nb) partner. Biologically active peptides are often fused with larger carrier proteins with the goal of modifying pharmacokinetic properties.^27^ Secondly, we examined whether there are advantages in terms of agonist activity or pathway selectivity when the ligand is linked to a nanobody that specifically binds to NK1R, as compared to a control Nb lacking such binding capacity. Previous studies involving PTHR1 ligands have demonstrated enhanced potency upon conjugation with Nbs that bind to the receptor.^8^ Whether this trend extended to other receptors was unknown prior to these studies. Lastly, we investigated the influence of the Nb epitope’s location on the observed signaling properties of Nb-ligand conjugates. By utilizing the triply tagged NK1R construct in our experimental design, we were able to directly probe this question for the first time.

Relevant to the questions above, the consequences of conjugating Nbs with ligands depended on the ligand used and the signaling pathway under study (**Figure 5)**. Conjugation of NKA and SP_6-11_ with Nbs caused consistently minor weakening in Gq signaling relative to comparator peptides **(Figure 2)**. In contrast, Nb-NKA and especially Nb-SP_6-11_, conjugates exhibited divergent properties for inducing cAMP accumulation and β-arrestin2 recruitment **(Figure 2)**. For G_s_ signaling, the conjugation of NKA with certain tag-binding Nbs (Nb_Alfa_ and Nb_6E_) resulted in highly potent partial agonists of cAMP signaling, whereas other Nb-NKA conjugates exhibited weak activity (**Figure 2**). In β-arrestin2 recruitment assays, all Nb-NKA and especially Nb-SP_6-11_ conjugates displayed reduced potency, which was not dependent on the identity of the Nb (binding vs. non-binding) used **(Figure 2)**.

**Figure 5.**
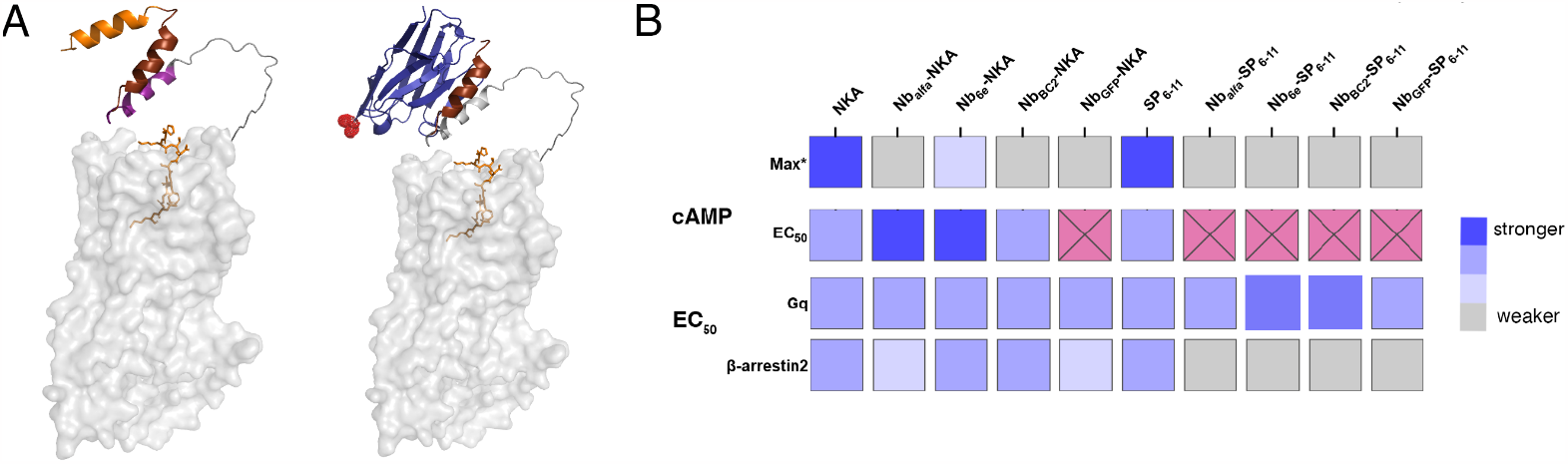
Model of epitope-tagged NK1R structures generated using AlphaFold 2 and summary of conjugate signaling activities. **(A)** In the model in the left, the epitope tags are highlighted as follows: alfa orange, 6e maroon and BC2 purple. Compound bound to the receptor is NKA shown as orange sticks with receptor in gray. In the model in the right Nb_6e_ is represented in blue and red spheres mark the *C*-terminus, which serves as the site of peptide conjugation. **(B)** Heat map summarizing ligand potencies (relative EC_50_ values) and efficacies (normalized maximal response values) for cAMP signaling and ligand potencies for G_q_ and β-arrestin2 signaling. Pink squares with X represent correspond to compounds for which activity was too weak to accurately calculate an EC_50_. *Max activity values were calculated by normalizing the response at 1 μM for conjugates to that of G3NKA or G3SP_6-11_ at 10 μM

Several other Nb-ligand conjugate properties, including performance in a competition binding assay, durability of G_s_ signaling responses and transcriptional outputs are affected by Nb identity. Nb_alfa_-NKA and Nb_6e_-NKA induced sustained cAMP production on cells expressing tag-NK1R but not WT-NK1R, highlighting the role of Nb binding (**Figure 3 & 4)**. Since Nb and NKA bind to distinct sites, a bitopic mode of interaction may contribute to the prolonged signaling observed. Complete ligand release from the receptor necessitates dissociation at both binding sites. Supporting this hypothesis, the addition of the NK1R antagonist, spantide I (1 μM),^26^ did not abolish the sustained response (**Figure S10**). Prolonged signaling responses have also been observed upon covalent tethering of a GPCR ligand to its receptor.^28^ Compounds with a long duration of action are useful in a variety of contexts^29^, such as long-acting PTHR1 agonists which are used to treat hypoparathyroidism.^29,30^ The findings here encourage exploration of Nb conjugation to facilitate the rational design of long-lasting ligands.

Structural modeling can provide a window for hypothesis generation related to conjugate signaling. To facilitate visualization of a hypothetical complex between a Nb-ligand conjugate and tagged-NK1R, we generated a model using the output of Alphafold2^31^ (**Figure 5A**). This model highlights how the different locations of epitope tags within tagged NK1R could lead to variation in the conjugate-receptor complex geometry when using Nbs that bind to these sites. Past work has characterized interactions between peptide ligands and NK1R that play a crucial role in receptor activation.^22^ In these previous studies, strong G_s_ signaling required interactions between the ligand and the extracellular loops of NK1R, whereas G_q_ signaling did not.^22^ We hypothesize that the geometric constraints imposed by Nb binding at different sites might dictate differences in the mode of receptor engagement for ligands linked to Nbs. We speculate that benefits imparted by ligand tethering are balanced against detrimental alterations in the mode of ligand-receptor interactions caused by steric clashes between Nb and receptor extracellular loops.

Some conjugates showed unexpectedly strong G_s_ and G_q_-driven transcriptional outputs, particularly in the case of Nb-NKA conjugates. **(Figure 4)**. GPCRs influence cellular functioning in large part through transcriptional outputs, which has been shown to be related to the subcellular localization of signaling.^32^ While GPCR activation can trigger immediate and transient cellular responses via signaling involving intracellular second messengers, the process of transcription is more intricate, complex, and slower in nature. We hypothesize that the enhanced G_s_ and G_q_-driven transcription observed might be mechanistically related to the prolonged cAMP responses observed.

## Conclusions

We have demonstrated that the conjugation of peptide agonists of GPCR signaling with Nbs that bind to the same receptor can provide compounds with altered signaling profiles relative to the prototype agonists. Interestingly, although the conjugation reduced the activation of certain signaling pathways, it significantly improved the transcriptional response of Nb-NKA conjugates relative to conventional ligands. This trend may relate to the prolonged or enhanced signaling observed for these conjugates in certain pathways. Our findings also emphasize the importance of carefully considering the location of epitopes used for ligand tethering, as the Nb binding site has a notable impact on the agonistic behavior of the conjugates. Given that all epitope tags used in this study were inserted consecutively, it seems likely that small variations in tethering agent binding site and orientation, such as those accessible from evaluating multiple target binding Nb/Ab clones, can play an important role in determining conjugate bioactivity profiles. These findings lay the groundwork for application of Nbs that recognize receptor epitopes outside the conserved ligand-binding pocket of GPCRs to generate conjugates with useful properties such as prolonged duration of action and high specificity.

## Methods

### Solid-phase peptide synthesis

Peptides were synthesized on a Gyros Protein Technologies PurePep Chorus automated peptide synthesizer (Uppsala, Sweden) by Fmoc-SPPS on a 0.05 mmol scale using Rink amide resin (ChemPep, 0.51 mmol/g). Fmoc deprotection was achieved using 20% piperidine/N,N-dimethylformamide DMF (v/v). Couplings were carried out in DMF using 8 equivalents relative to the resin loading of Fmoc–amino acid acid/PyAOP/ N,N-diisopropylethylamine (DIPEA) (1:1:2 molar ratio). Peptide cleavage from the resin and removal of side-chain protecting groups was achieved using 90% trifluoroacetic acid (TFA)/5% tri-isopropylsilane (TIPS) /5% H_2_O for 2 h at 25 °C. The peptides were precipitated with chilled diethyl ether, pelleted by centrifugation (3,000 RPM for 5 minutes) and then lyophilized in 50% acetonitrile (ACN)/0.1% TFA/H_2_O.

### Reversed-phase high-performance liquid chromatography (RP-HPLC) and LC–MS

Peptides were purified using a preparative C_18_ column (Aeris PEPTIDE 5 μm XB-C18, LC Column 250 x 21.2 mm, AXIA™ Packed, Phenomenex, flow rate 10 mL/min) in a Shimadzu LC-20AR solvent delivery system with a gradient of 20–70% B over 25 min. Solvents consisted of 0.1% TFA in H_2_O (solvent A) and 0.1% TFA in ACN (solvent B). The molecular weight of the fractions collected was analyzed on an ESI-MS. Fractions with the desired mass were further analyzed for purity by analytical reversed-phase (RP) high-performance liquid chromatography (HPLC) and lyophilized.

Analytical RP-HPLC on an analytical C_18_ column (Aeris 5 μm PEPTIDE XB-C18, 4.6 × 250 mm, 100 Å, Phenomenex, flow rate 1 mL/min) connected to a Shimadzu LC-40D solvent delivery system equipped with a SIL-40C autoinjector and an SPD-40D UV–vis detector was used to determine the purity of purified peptides. A linear gradient of 0–50% B over 50 min was used, and absorbance data were collected at 214 nm.

Mass spectrometry data were acquired on a Waters Xevo qTOF LC/MS. Samples were resolved by RP-HPLC on a Hamilton PRP-h5 column (5 μM particle size, 300 Å pore size) and analyzed in positive ion mode. Data acquisition and processing were carried out using MassLynx software.

### Nanobody recombinant expression

BL21(DE3) *E. coli* were heat shock transfected with pET26b(+) plasmids encoding corresponding nanobodies and cultured in Terrific Broth medium containing ampicillin (100 g/mL) or kanamycin (50 g/mL). Transformed bacteria were employed to create a preculture, which was then used to inoculate a full-size culture (1-4 L), which was shaken at 37 °C until mid-log phase (optical density at 600 nm between 0.6 and 0.8). Protein expression was induced by the addition of isopropyl-d-1-thiogalactopyranoside (IPTG, 1 mM) and the induced culture was shaken at 30°C overnight. Bacteria were harvested by centrifugation at 6,000 RPM (Avanti J Series centrifuge) for 20 minutes and suspended in NTA wash buffer (tris buffered saline + 10 mM imidazole, pH 7.5) containing lysozyme and incubated on ice for 10 min. The cells were then lysed using sonication (3x), and the lysate was centrifuged at 16,000 RPM for 40 minutes to pellet lysate.

The supernatant was then passed through a fritted column containing nickel NTA beads (His Pur^TM^ Ni-NTA Resin) equilibrated with nickel NTA wash buffer (tris-buffered saline (TBS) + 10 mM imidazole, pH 7.5). Following the initial flowthrough, the beads were washed three times with a Nickel NTA wash buffer. Then, 10 mL of Nickel NTA elution buffer (TBS + 150 mM imidazole, pH 7.5) was used to elute the bound protein. Size exclusion chromatography was performed on the sample (HiLoad TM 16/600 Superdex 200 pg column, Cytiva Akta^TM^ / Pure) with an isocratic gradient of TBS (flow rate 1 mL/min). Spin filtration columns (Amicon Ultra-15, regenerated cellulose, 10 kDa nominal molecular weight limit) were used to concentrate fractions containing the protein of interest. Protein concentrations were measured by UV spectroscopy measuring absorption at 280 nm using a Nanodrop spectrometer (Thermo Scientific).

Plasmid and nanobody sequences are provided in the SI

### Nanobody conjugation via sortagging

Sortagging reactions were carried out in sortase buffer (10 mM CaCl_2_, 50 mM Tris, 150 mM NaCl, pH 7.5) containing the nanobody bearing a sortase recognition motif (LPETGG) (20-200 μM), triglycine functionalized peptide (500-1000 μM), and Sortase-A 5M (10-20 μM). After incubation at 12 °C with agitation for 16 h, the reaction was incubated with nickel NTA beads to capture Sortase-A 5M and unreacted nanobody. Uncaptured material was further purified using disposable desalting columns to remove triglycine-peptides (Cytiva PD-10 Sephadex^TM^ G-25M). Fractions containing the product were combined and then concentrated by spin filtration (Amicon Ultra 0.5 mL Centrifugal Filters 10 kDa NMWL). The molecular weight of the conjugates was confirmed by LC-MS.

### Generation of stable cell lines

Human embryonic kidney 293 (HEK293) cells stably expressing a cAMP-responsive luciferase^33^ were transfected (Lipofectamine™ 3000 Transfection Reagent) with either NK1R wide-type or epitope-tagged NK1R (plasmids sequences provided in the SI). After 24 hours, cells were cultured with Geneticin™ Selective Antibiotic (G418 sulfate; 1 mg/mL). Clonal lines were isolated by limiting dilution to provide a cell line that stably expresses cAMP biosensor and receptor of interest that can be grown without selection antibiotic. Cells were routinely cultured in Dulbecco’s modified Eagle’s medium (DMEM) supplemented with 10% fetal bovine serum (FBS) and 1x penicillin/streptomycin at 37 °C in 5% CO_2_.

### Flow cytometry analyses

Suspensions of cells in phosphate-buffered saline (PBS) containing 2% bovine serum albumin (BSA) (w/v) (PBS/BSA) were incubated on ice with nanobodies functionalized with biotin at various concentrations (0.1 pM – 1 μM) for 20 min. Cells were pelleted by centrifugation (500 RPM for 3 min), washed with PBS/BSA (3x), then resuspended in PBS/BSA containing APC-streptavidin (BioLegend, 405207) diluted 1:2000 for 30 min, pelleted and washed (3x) a second-time prior flow cytometry analysis on a CytoFlex flow cytometer (Beckman Coulter). Cells were first gated based on forward and side scatter to select intact cells. Data were analyzed using FlowJo. Nanobody binding concentration-response curves were generated using the median fluorescence intensity (MFI) of labeled cells. For the competition binding assay, cells were incubated on ice with increasing concentrations of unlabeled conjugates or free peptides in presence of SP-AF647 (30 nM) for 15 min. For the control samples, cells were incubated with SP-AF647 alone. Then cells were washed with PBS/BSA (3x) and resuspended in PBS/BSA analyzed as described above. Mean fluorescence intensity (MFI) values in the FL4 channel were determined for each sample. Percent inhibition was calculated through normalization of MFI in the absence of inhibitor to MFI observed in the presence of inhibitor.

### Measurement of cAMP response

HEK-293 stably expressing the Glosensor cAMP reporter (Promega Corp.) ^21^ and NK1R wide-type or epitope-tagged NK1R were seeded in a white-walled 96-well flat clear bottom plate (Corning #3610) and grown to confluency at 37°C in a 5% humidified CO_2_ incubator. The growth medium was removed from confluent monolayers of cells and replaced with 90 μL CO_2_ independent medium containing D-luciferin (0.5 mM) until a stable baseline level of luminescence was established (12 min). Compounds were added at various concentration (1 pM – 10 μM), and the luminescence response was measured every 2 minutes for 12 minutes using a Biotek Neo2 plate reader (ligand on). Concentration-response curves were generated using the maximal luminescence response (6-8 minutes) after ligands addition using GraphPad Prism (v.9.0) software. For the measurement of cAMP signaling duration experiments, cells were incubated with the compounds at various concentrations for 12 min. Then, the medium was removed along with unbound ligand. Fresh CO_2_ independent medium containing D-luciferin (0.5 mM) with or without spantide I (1 μM) was added and the luminescence response was measured for 30 min using a Biotek Neo2 plate reader (washout). GraphPad Prism software (v.9.0) was used to calculate the area under the curve (AUC) from the kinetic data.

### Bioluminescence Resonance Energy Transfer (BRET) Assay

Cells stably expressing epitope-tagged NK1R grown in a 6-well plate were transiently transfected (Lipofectamine™ 3000 Transfection Reagent) with 36 ng of β-arrestin2-RLucII and 504 ng of the acceptor protein (rGFP-CAAX or rGFP-FYVE) at a 14:1 (w/w) ratio, seeded in a white-walled 96-well plate flat clear bottom (Corning #3610) and grown to confluency at 37°C in a 5% humidified CO_2_ incubator. Growth medium was removed from confluent monolayers of cells, and 100 μL of the compounds at various concentrations (0.1 pM -1 μM) diluted in 5 mM HEPES + 1x HBSS + prolume purple (1 μM) was added. BRET signal was measured every 2.5 minutes for 30 minutes using a Biotek Neo2 plate reader by measuring luminescence at 515 and 410 nm and calculating the emission ratio for 515/410 nm.

### Bioluminescence Resonance Energy Transfer (BRET) TRUPATH assay

Cells stably expressing epitope-tagged NK1R were transiently transfected (Lipofectamine™ 3000 Transfection Reagent) with 1 μg of Gq TRUPATH G?/?/? biosensor plasmid24 seeded in a white-walled 96-well flat clear bottom plate (Corning #3610) and grown to confluency at 37°C in a 5% humidified CO_2_ incubator. Growth medium was removed from confluent monolayers of cells, and 100 μL of the compounds at various concentrations (0.1 pM -1 μM) diluted in 5 mM HEPES + 1x HBSS + methoxy e-Coelenterazine (prolume purple) (1 μM) was added. BRET signal was measured every 5 minutes for 40 minutes using a Biotek Neo2 plate reader by measuring luminescence at 515 and 410 nm and calculating the emission ratio for 515/410 nm.

### Intracellular Ca^2+^ mobilization signaling assay

HEK-293 stably expressing the Glosensor cAMP reporter (Promega Corp.)^21^ and epitope-tagged NK1R were seeded in a black-walled 96-well plate flat clear bottom (Corning) and grown to confluency at 37°C in a 5% humidified CO_2_ incubator. The cells were loaded with the FLIPR Calcium-6 no-wash dye (Product # R8190) dissolved in 20 mM HEPES buffer + 1x Hanks’ Balanced Salt Solution (HBSS), 2.5 mM probenocid, pH 7.4 and incubated for 2 hours at 37°C in 5% CO_2_. Intracellular Ca^2+^ responses were evaluated in a Fluorometric Imaging Plate Reader (FLIPR; Molecular Devices, Sunnyvale, CA) using a cooled CCD camera with excitation at 470-495 nM and emission at 515-575 nM in response to 1 μM of the compounds. Each plate’s camera gain and intensity were adjusted to produce a minimum of 1000 arbitrary fluorescence units (AFU) baseline fluorescence. A baseline fluorescence reading was taken prior to the addition of the compounds, followed by fluorescent readings every second for 300 seconds. The Delta F/F_0_ value was calculated, where F_0_ is the baseline level of fluorescence and Delta F is the change in fluorescence from the baseline level.

### Transcriptional reporter assay

Cells stably expressing epitope-tagged NK1R were transiently transfected (Lipofectamine™ 3000 Transfection Reagent) with 1 μg of pGL4.29[*luc2P*/CRE/Hygro] or pGL4.30[*luc2P*/NFAT-RE/Hygro] plasmids (Promega Corp.), seeded in a white-walled 96-well flat clear bottom plate (Corning #3610), and grown to confluency at 37°^34^C in a 5% humidified CO_2_ incubator for 24h. Cells were then incubated with ∼35 nM of the compounds for 17 hours. Transcription was measured using the Bright-Glo^TM^ Luciferase Assay System (Promega Corp.) according to manufacture protocol with small modifications. Briefly, 30 μl of Bright-Glo^TM^ was added to the cells and luminescence response was measured using a Biotek Neo2 plate reader. Fold induction was calculated by dividing the relative luminescence recorded for induced cells by the relative luminescence of control cells. Normalized values were then normalized again to index peptides (G3NKA or G3SP_6-11_).

### Computational Protein Modeling

The online tool ColabFold implementing Alphafold2^31, 34^ was used to predict tagged NK1R and Nb_6e_ structures.

## Acknowledgments

The authors B. Roth for providing the TRUPATH vector (Addgene Kit # 1000000163) used in this study. We acknowledge the NIDDK mass spectrometry core (J. Lloyd) for assistance. We acknowledge T. Gardella (Massachusetts General Hospital) for provision of HEK293 cells stably transfected with Glosensor cAMP reporter. We acknowledge L. Wingler (Duke) for helpful discussions. We acknowledge M. Bouvier (University of Montreal) for provision of plasmids used for β-Arrestin recruitment assays. This work was supported by the NIH Intramural Research Program (NIDDK) and by funding from the NIH Director’s Award (1ZIADK075157-02). Some figures were created with BioRender.com.

