## Supplementary material for "Semi-synthetic nanobody-ligand conjugates exhibit tunable signaling properties and enhanced transcriptional outputs at neurokinin receptor-1": SI

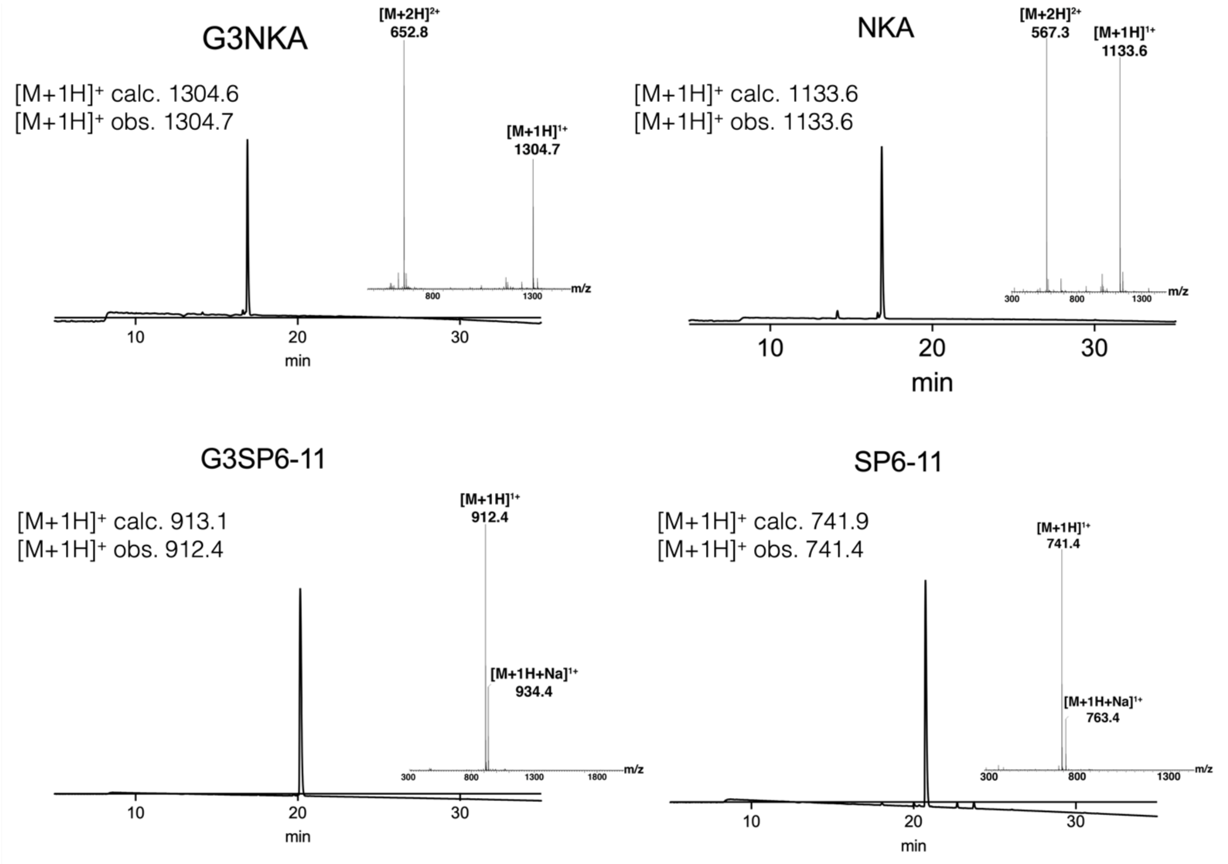


**Figure S1. Analytical HPLC and high-resolution MS analysis of G3NKA, NKA, G3SP$_{6\text{-}11}$ and SP$_{6\text{-}11}$.** HPLC and mass spectrometry were performed according to Methods.

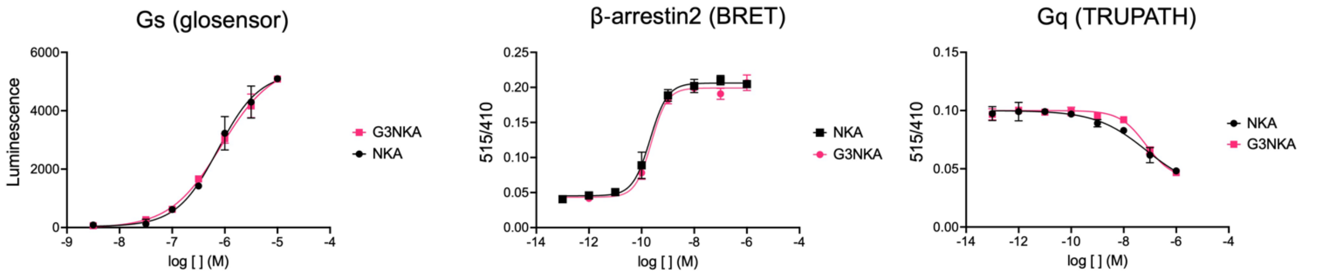


| $EC_{50}$ | NKA | G3NKA |
|---|---|---|
| Mean (nM) | 384.2 | 604.7 |
| SEM (±) | 183.4 | 191.2 |

| $EC_{50}$ | NKA | G3NKA |
|---|---|---|
| Mean (nM) | 3.5 | 0.5 |
| SEM (±) | 2.1 | 0.3 |

| $EC_{50}$ | NKA | G3NKA |
|---|---|---|
| Mean (nM) | 32.6 | 40.8 |
| SEM (±) | 14.9 | 21.3 |

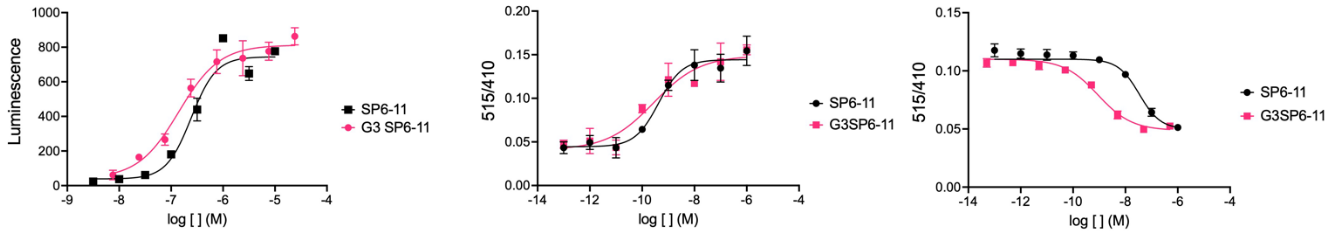


| $EC_{50}$ | SP6-11 | G3SP6-11 |
|---|---|---|
| Mean (nM) | 231.1 | 102.5 |
| SEM (±) | 71.5 | 20.6 |

| $EC_{50}$ | SP6-11 | G3SP6-11 |
|---|---|---|
| Mean (nM) | 0.4 | 0.2 |
| SEM (±) | 0.1 | 0.1 |

| $EC_{50}$ | SP6-11 | G3SP6-11 |
|---|---|---|
| Mean (nM) | 27.5 | 0.3 |
| SEM (±) | 14.3 | 0.2 |

**Figure S2. Comparison of the activity NKA and $SP_{6-11}$ to their triglycine analogues for inducing Gs, β-arrestin2 and $G_q$ responses.** Representative concentration-response curves (mean ± SD from technical replicates). $EC_{50}$ was calculated from the fitting of a sigmoidal concentration-response model to data from three or more independent experiments using GraphPad Prism.

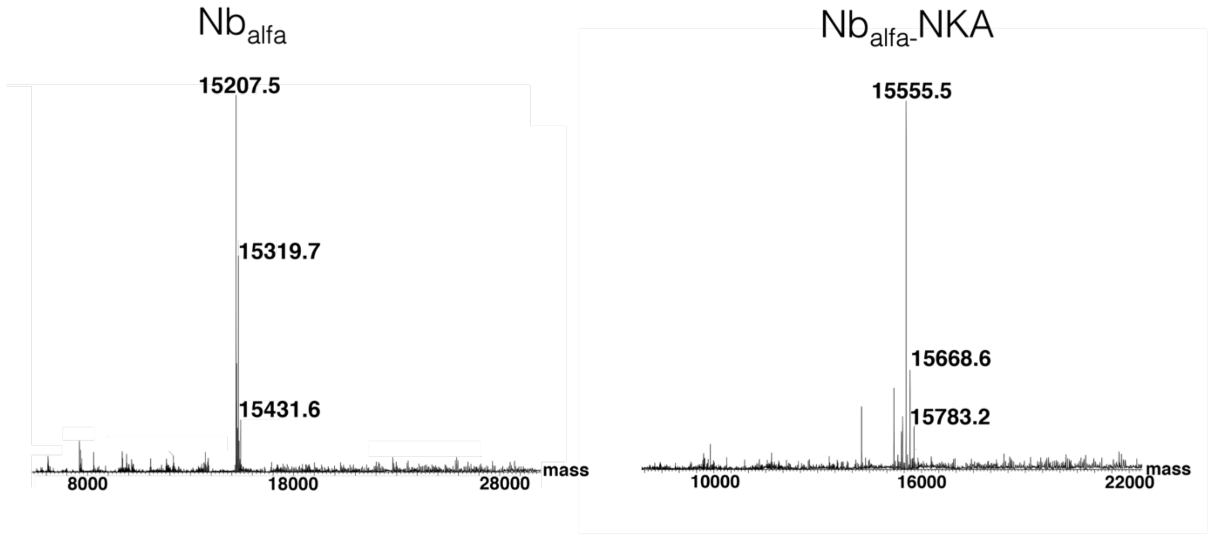

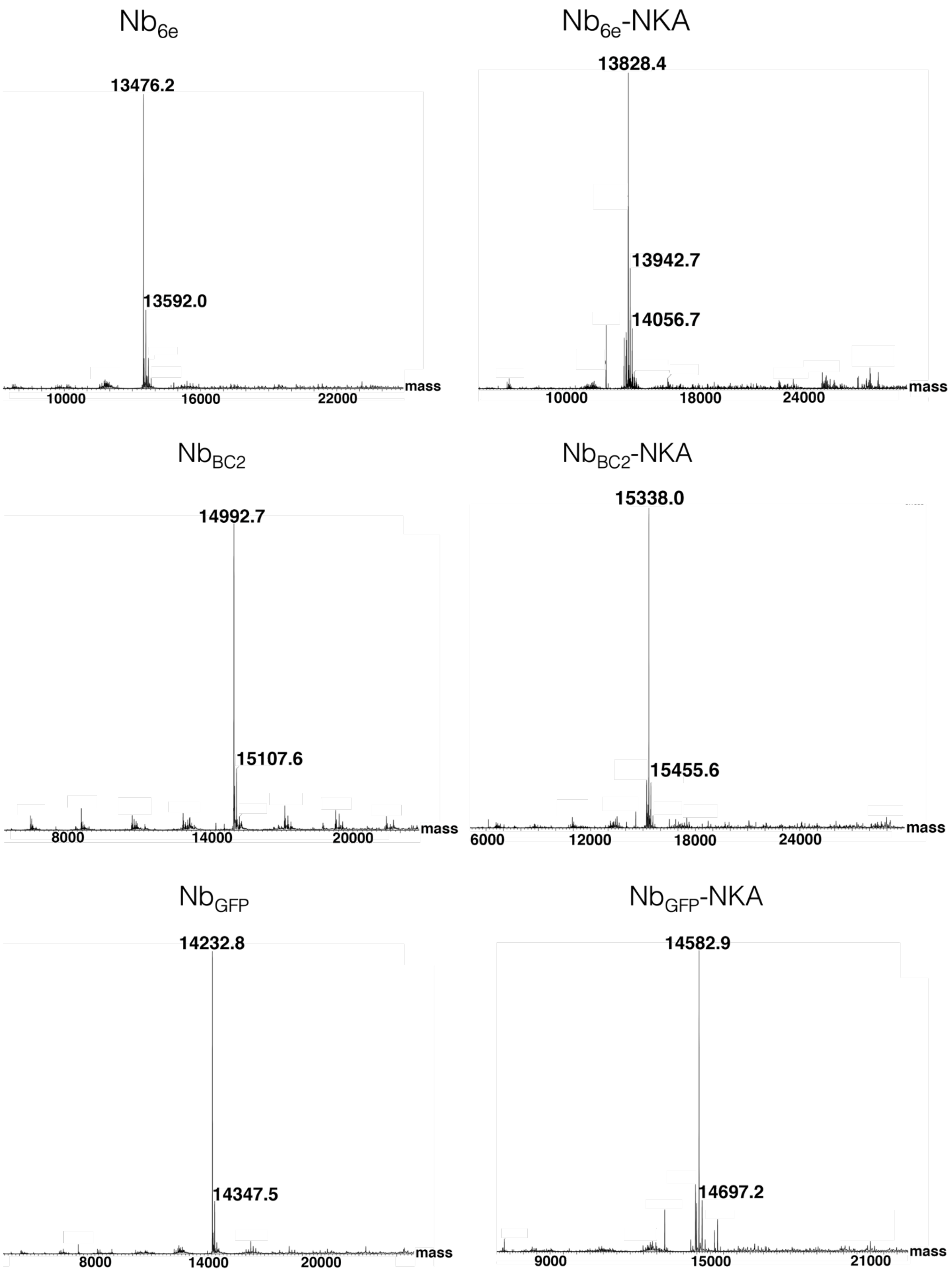

Nb6e
13476.2
13592.0
10000
16000
22000
mass
Nb6e-NKA
13828.4
13942.7
14056.7
10000
18000
24000
mass
NbBC2
14992.7
15107.6
8000
14000
20000
mass
NbBC2-NKA
15338.0
15455.6
6000
12000
18000
24000
mass
NbGFP
14232.8
14347.5
8000
14000
20000
mass
NbGFP-NKA
14582.9
14697.2
9000
15000
21000
mass

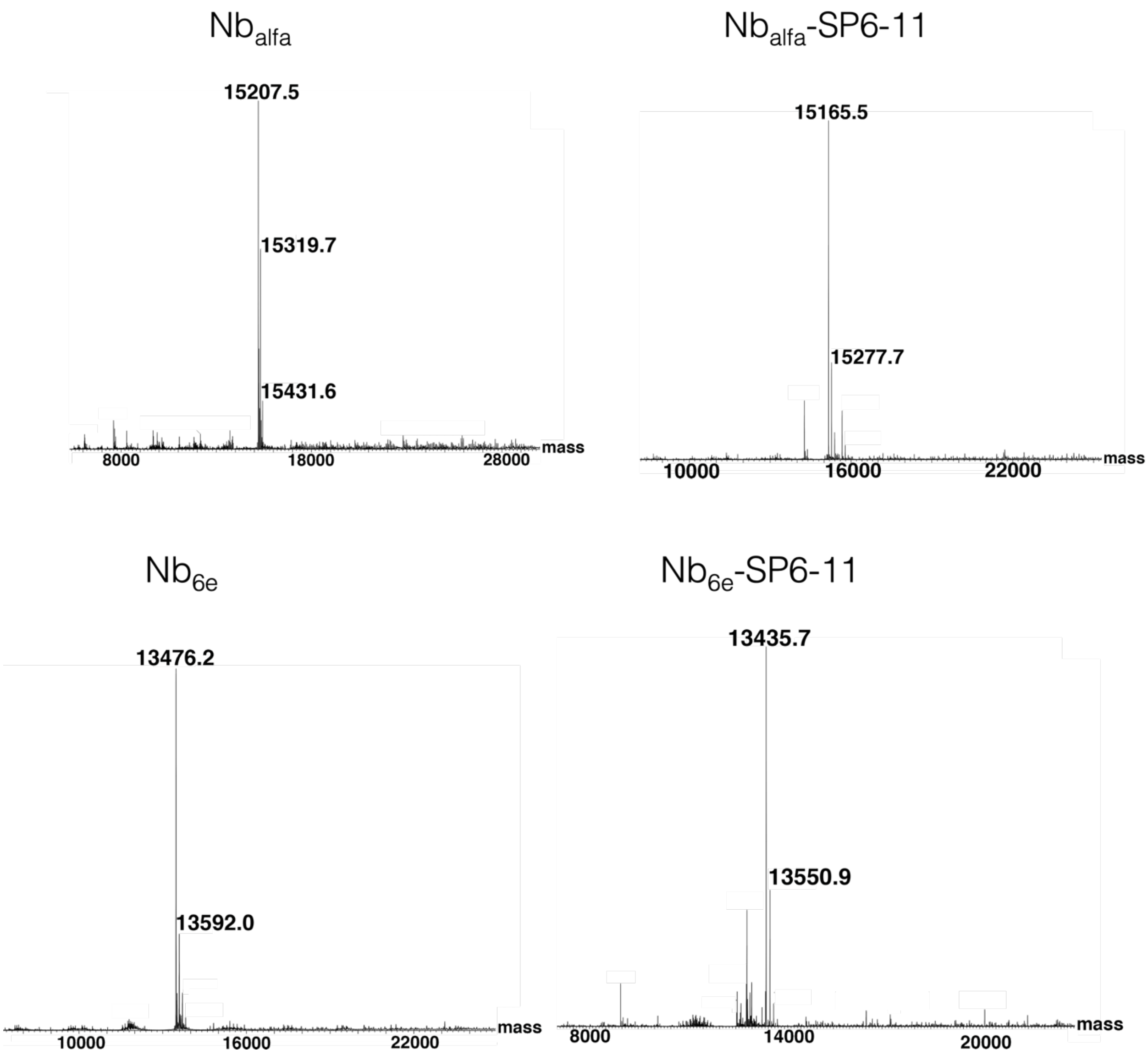

Nb$_{alfa}$
15207.5
15319.7
15431.6
8000
18000
28000
mass
Nb$_{alfa}$-SP6-11
15165.5
15277.7
10000
16000
22000
mass
Nb$_{6e}$
13476.2
13592.0
10000
16000
22000
mass
Nb$_{6e}$-SP6-11
13435.7
13550.9
8000
14000
20000
mass

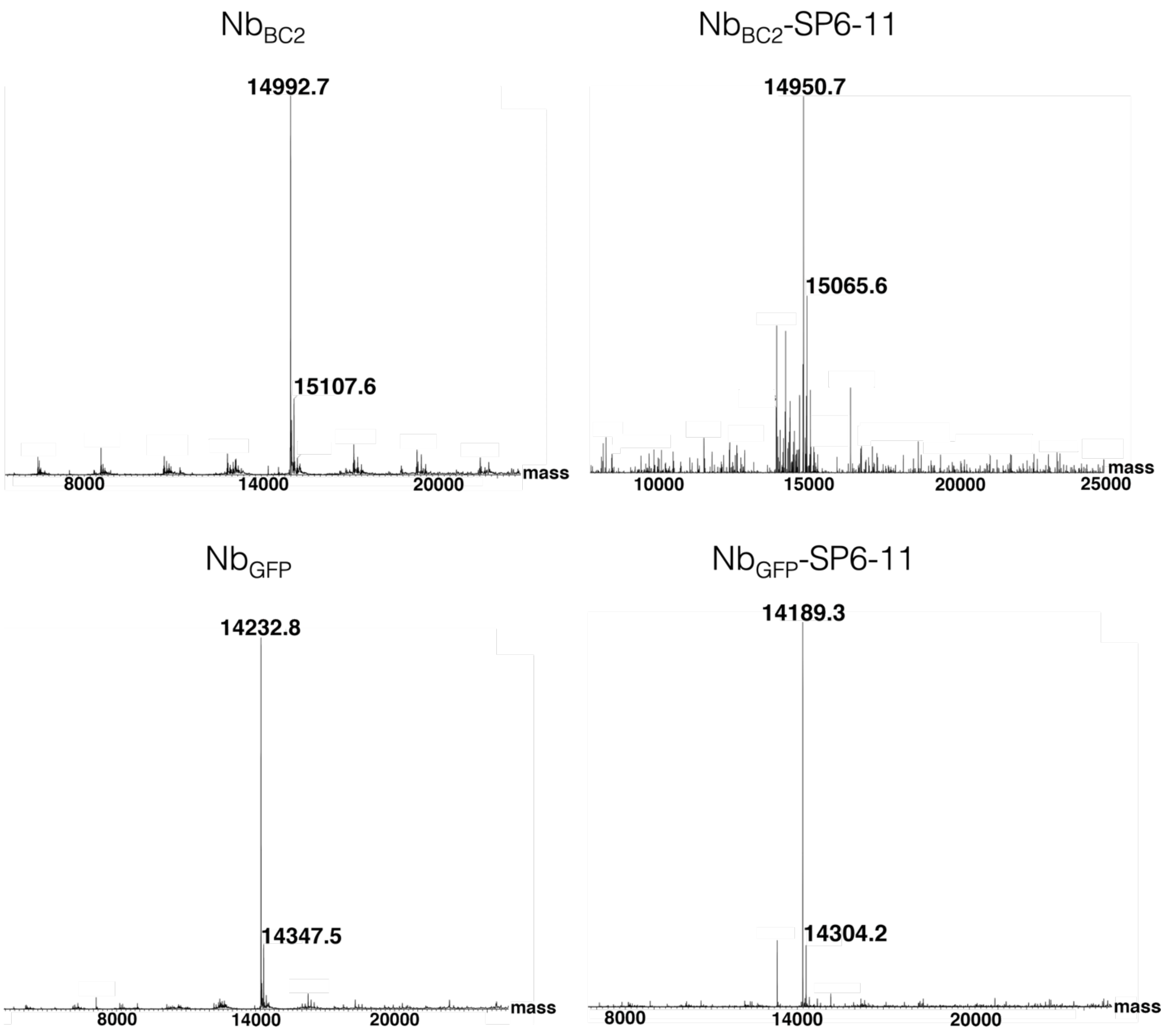


**Figure S3. MS of the nanobodies before and after conjugation to G3NKA or G3SP$_{6-11}$.** Mass spectra and deconvoluted masses were acquired according to Methods.

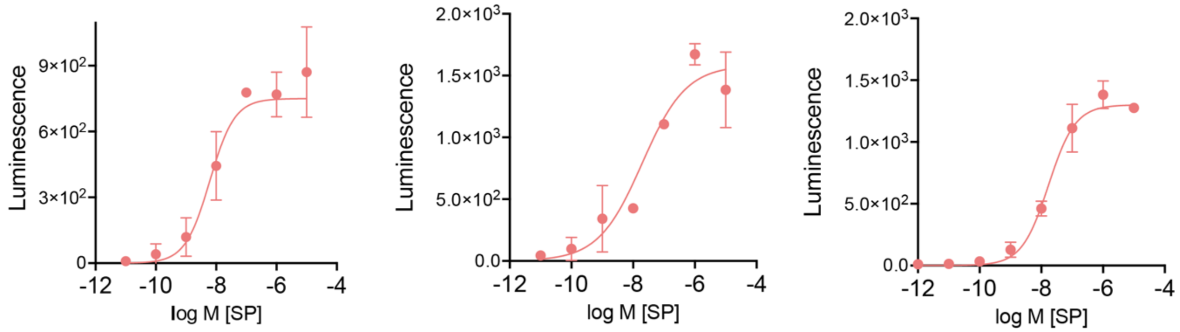


**Figure S4. Replicate experiments of substance P (SP) activation of epitope-tagged NK1R via Glosensor assay.** Data points correspond to mean ± SD from technical replicates.

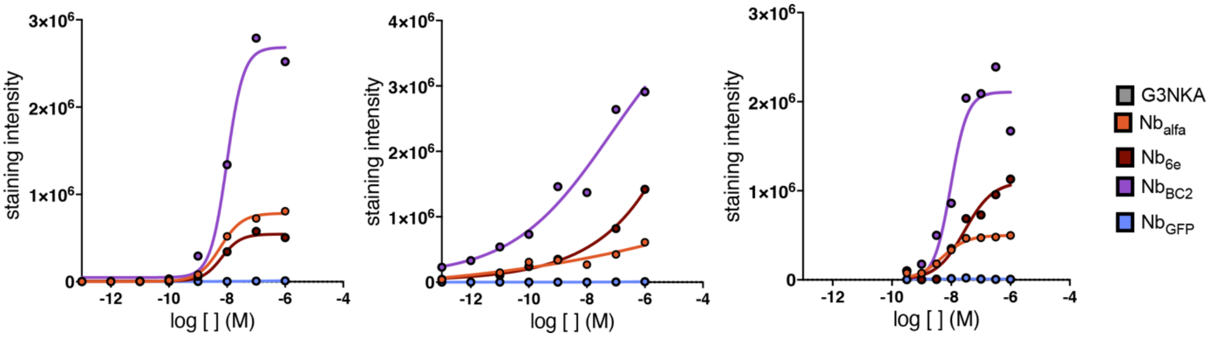


**Figure S5. Independent replicates for flow cytometry analysis of biotin-labeled nanobodies binding to epitope-tagged NK1R cells.** Staining and analysis was performed according to Methods.

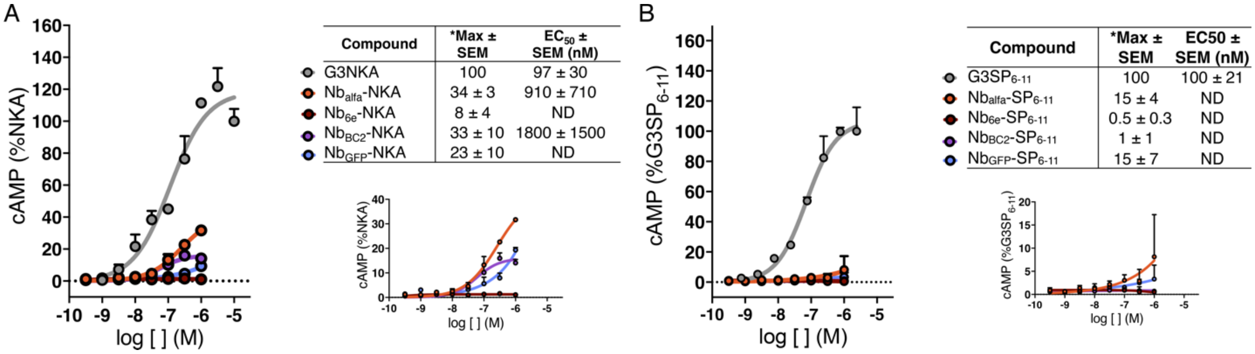


| Compound | *Max ± SEM | EC50 ± SEM (nM) |
|---|---|---|
| G3NKA | 100 | 97 ± 30 |
| $Nb_{alfa}$-NKA | 34 ± 3 | 910 ± 710 |
| $Nb_{6e}$-NKA | 8 ± 4 | ND |
| $Nb_{BC2}$-NKA | 33 ± 10 | 1800 ± 1500 |
| $Nb_{GFP}$-NKA | 23 ± 10 | ND |

| Compound | *Max ± SEM | EC50 ± SEM (nM) |
|---|---|---|
| $G3SP_{6-11}$ | 100 | 100 ± 21 |
| $Nb_{alfa}$-$SP_{6-11}$ | 15 ± 4 | ND |
| $Nb_{6e}$-$SP_{6-11}$ | 0.5 ± 0.3 | ND |
| $Nb_{BC2}$-$SP_{6-11}$ | 1 ± 1 | ND |
| $Nb_{GFP}$-$SP_{6-11}$ | 15 ± 7 | ND |

**Figure S6. Effects of peptides and conjugates on cAMP production in NK1R wild-type cells.** HEK293 cells stably transfected with Glosensor cAMP reporter (Promega Corp.)[23] and wild-type NK1R were treated with varying concentrations (1 pM – 10 mM) of the indicated peptides or conjugates. Activation was assessed by cAMP production after 6 min (n = 3) as described in Methods. (**A, B**) Representative concentration-response curves (mean ± SD). Insets of concentration-responses without G3-peptides included are shown (under table) to allow better visualization of the activity of the weaker compounds. Curves result from the fitting of a sigmoidal concentration-response model to data. *Max activity values were calculated by normalizing the response at 1 mM for conjugates to that of G3NKA or $G3SP_{6-11}$ at 10 mM (n = 3). $EC_{50}$ for conjugates with maximal responses lower than 30% were not determined (ND).

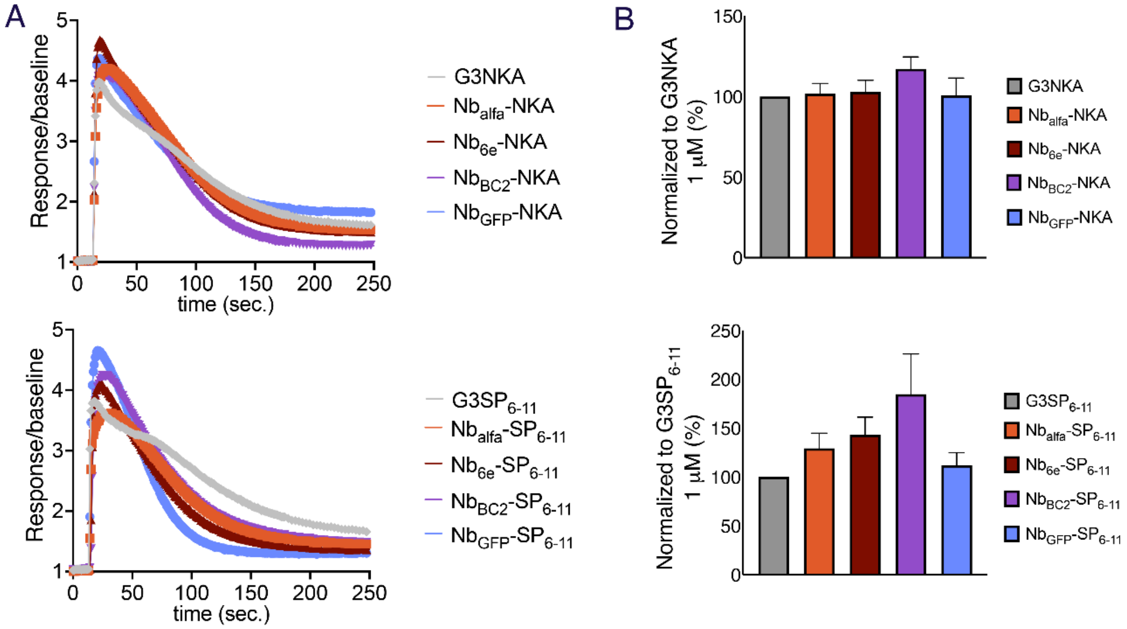


**Figure S7. Effects of peptides and conjugates on intracellular $Ca^{2+}$ mobilization**. HEK293 cells stably expressing tagged NK1R were treated the indicated peptides or conjugates. **A.** Representative FLIPR traces of the fluorescenc over the basal fluorescence following the addition of the compounds at 1 μM to the cells. **B.** Bar graphs represent meai from three independent biological replicates. Data were normalized to G3NKA or $G3SP_{6-11}$ responses at 1 μM.

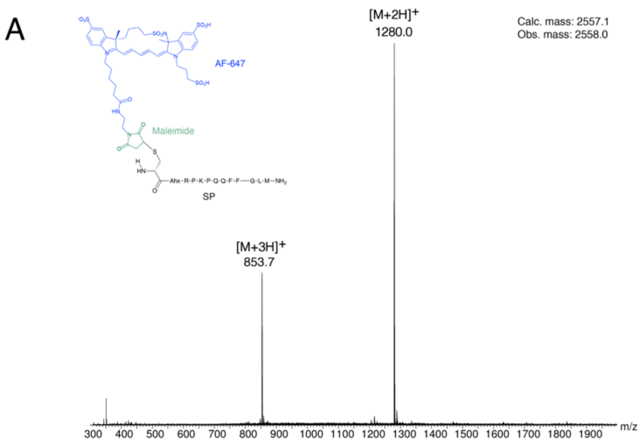


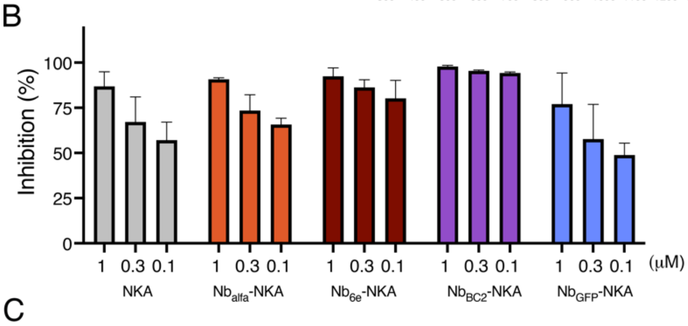


| | | 1 uM | 300 nM | 100 nM |
|---|---|---|---|---|
| % inhibition ± SEM | NKA | 87 ± 5 | 67 ± 8 | 57 ± 6 |
| | $Nb_{alfa}$-NKA | 91 ± 1 | 74 ± 6 | 66 ± 2 |
| | $Nb_{6e}$-NKA | 92 ± 3 | 86 ± 2 (*) | 80 ± 6 (**) |
| | $Nb_{BC2}$-NKA | 98 ± 0 | 95 ± 0 (***) | 94 ± 0 (****) |
| | $Nb_{GFP}$-NKA | 77 ± 10 | 58 ± 14 | 49 ± 4 |

( ) Dunnett's multiple comparison test vs NKA

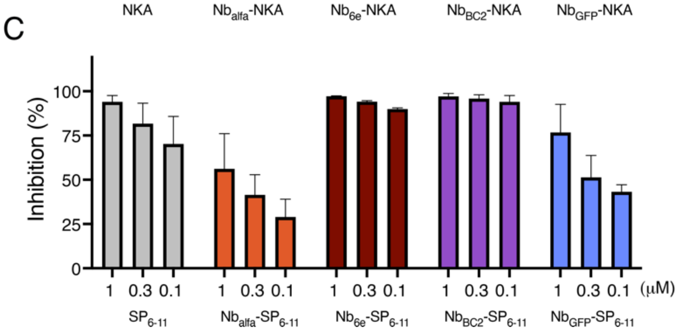


| | | 1 uM | 300 nM | 100 nM |
|---|---|---|---|---|
| % inhibition ± SEM | $SP_{6-11}$ | 94 ± 2 | 82 ± 7 | 70 ± 9 |
| | $Nb_{alfa}$-$SP_{6-11}$ | 56 ± 11 (****) | 41 ± 7 (****) | 29 ± 6 (****) |
| | $Nb_{6e}$-$SP_{6-11}$ | 97 ± 0 | 94 ± 0 | 90 ± 1 |
| | $Nb_{BC2}$-$SP_{6-11}$ | 97 ± 1 | 96 ± 1 | 94 ± 2 (**) |
| | $Nb_{GFP}$-$SP_{6-11}$ | 77 ± 9 | 51 ± 7 (***) | 43 ± 2 (**) |

( ) Dunnett's multiple comparison test vs NKA

**Figure S8.** B**inding assay for measurement of Nb-ligand conjugate competition with labeled SP. (A)** Chemical structure and high-resolution MS analysis of SP-AF647. Binding assay was performed using flow cytometry to assess the ability of **(B)** NKA and Nb-NKA or **(C)** $SP_{6-11}$ and Nb-$SP_{6-11}$ to outcompete fluorescently labeled SP for binding to epitope-tagged NK1R. Values correspond to mean ± SEM from 3 biological replicates. Differences were evaluated for statistical significance with a one-way ANOVA followed by Dunnett correction. *$p < 0.03$, **$p < 0.002$, ***$p < 0.0002$, **** $p < 0.0001$

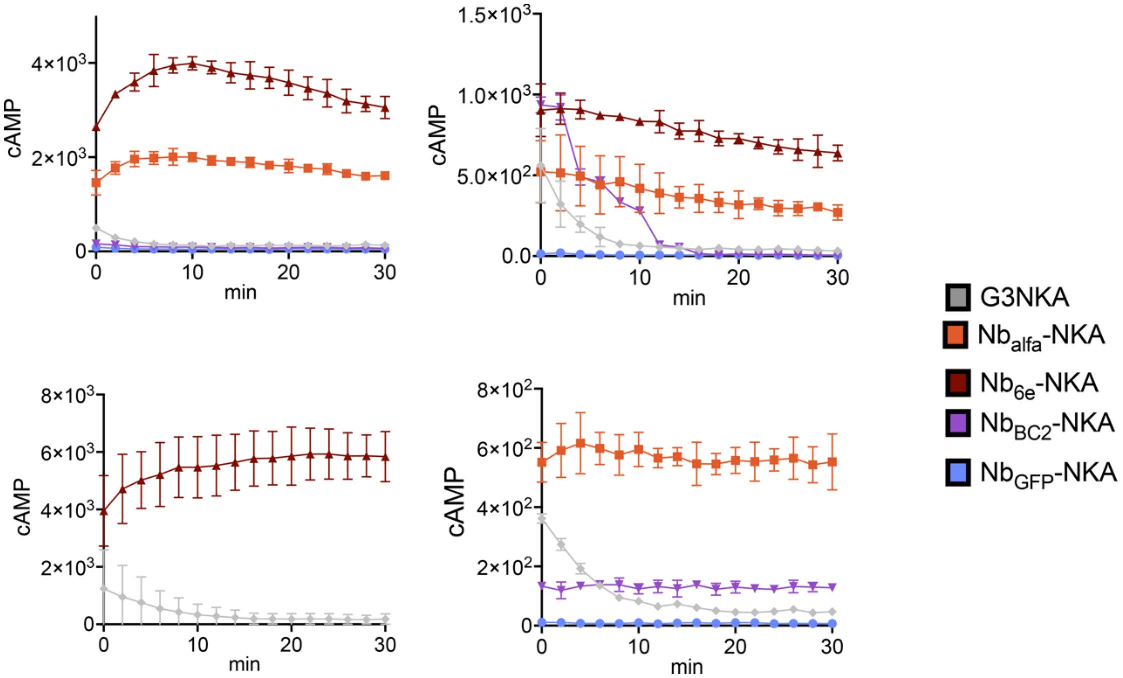


**Figure S9. Replicate experiments for measurement of the decay of cAMP production via washout assay in epitope-tagged NK1R.** Data points correspond to mean ± SD from technical replicates.

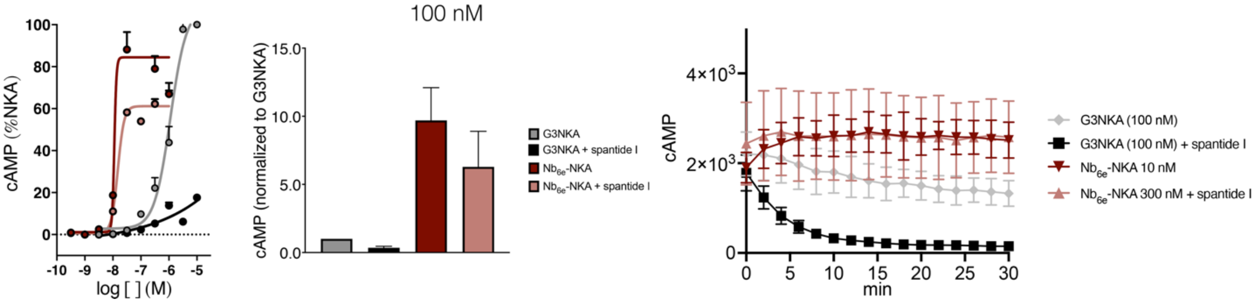


**Figure S10. Impact of spantide I on the dissipation of cAMP signaling (washout) for $Nb_{6e}$-NKA and G3NKA**. Excess/unbound ligand was removed, fresh media with 1 μM spantide I was added, and cAMP concentration was measured for an additional 30 min (washout) (n = 3). Washout assay data is summarized as the area under the curve (AUC) recorded for a 100 nM of peptide or conjugate. Data represent mean ± SEM from n=3 independent experiments. Representative concentration-response curves or cAMP production kinetics (mean ± SD) for washout assays.

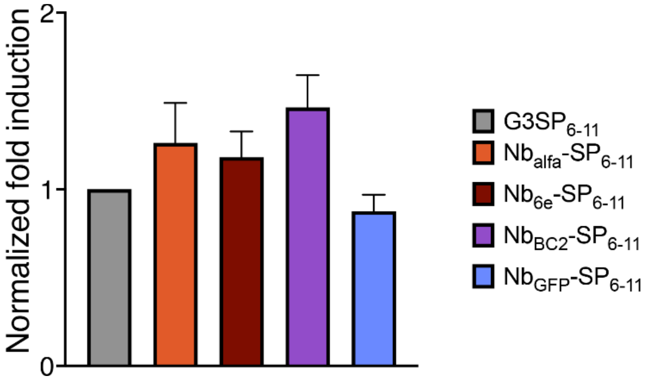


**Figure S11. Evaluation of G3SP6-11 peptide and conjugate performance on Gq transcription.** Transcriptional responses were evaluated through transfection of cells expressing epitope tagged NK1R with a luciferase reporter plasmid reporting on $G_q$ signaling. Cells were incubated with ~35 nM of the indicated peptides or conjugates for 17 hours and transcription was measured as described in Methods. Data represent mean ± SEM from n=3 independent experiments.

### Plasmids and nanobody sequences

Epitope-tagged NK1R (receptor sequence underlined)

gacggatcgggagatctcccgatcccctatggtgcactctcagtacaatctgctctgatgccgcatagttaagccagtatctgctccctgcttgtgtgt
tggaggtcgctgagtagtgcgcgagcaaaatttaagctacaacaaggcaaggcttgaccgacaattgcatgaagaatctgcttagggttaggcg
ttttgcgctgcttcgcgatgtacgggccagatatacgcgttgacattgattattgactagttattaatagtaatcaattacggggtcattagttcatagcc
catatatggagttccgcgttacataacttacggtaaatggcccgcctggctgaccgcccaacgacccccgcccattgacgtcaataatgacgtat
gttcccatagtaacgccaatagggactttccattgacgtcaatgggtggagtatttacggtaaactgcccacttggcagtacatcaagtgtatcatat
gccaagtacgccccctattgacgtcaatgacggtaaatggcccgcctggcattatgcccagtacatgaccttatgggactttcctacttggcagtac
atctacgtattagtcatcgctattaccatggtgatgcggttttggcagtacatcaatgggcgtggatagcggtttgactcacggggatttccaagtctcc
accccattgacgtcaatgggagtttgttttggcaccaaaatcaacgggactttccaaaatgtcgtaacaactccgccccattgacgcaaatgggcg
gtaggcgtgtacggtgggaggtctatataagcagagctctctggctaactagagaacccactgcttactggcttatcgaaattaatacgactcacta
tagggagacccaagctggctagcgtttaaacttaagcttggtaccgagctcggatccgccaccatgaagacaatcatcgccctgagctacatctt
ctgcctggtgttcgccggaccttctagactggaagaggagctgcgccggagactgaccgagcccggccaggccgaccaggaggccaagga
actggctagacagatcagcggccctgatagagtgcgggccgtgtcccactggagcagcatggataacgtcctcccggtggactcagacctctc
cccaaacatctccactaacacctcggaacccaatcagttcgtgcaaccagcctggcaaattgtcctttgggcagctgcctacacggtcattgtggt
gacctctgtggtgggcaacgtggtagtgatgtggatcatcttagcccacaaaagaatgaggacagtgacgaactattttctggtgaacctggcctt
cgcggaggcctccatggctgcattcaatacagtggtgaacttcacctatgctgtccacaacgaatggtactacggcctgttctactgcaagttccac
aacttctttcccatcgccgctgtcttcgccagtatctactccatgacggctgtggcctttgataggtacatggccatcatacatcccctccagccccgg
ctgtcagccacagccaccaaagtggtcatctgtgtcatctgggtcctggctctcctgctggccttcccccagggctactactcaaccacagagacc
atgcccagcagagtcgtgtgcatgatcgaatggccagagcatccgaacaagatttatgagaaagtgtaccacatctgtgtgactgtgctgatctac
ttcctcccccctgctggtgattggctatgcatacaccgtagtgggaatcacactatgggccagtgagatccccggggactcctctgaccgctaccac
gagcaagtctctgccaagcgcaaggtggtcaaaatgatgattgtcgtggtgtgcaccttcgccatctgctggctgcccttccacatcttcttcctcctg
ccctacatcaacccagatctctacctgaagaagtttatccagcaggtctacctggccatcatgtggctggccatgagctccaccatgtacaacccc
atcatctactgctgcctcaatgacaggttccgtctgggcttcaagcatgccttccggtgctgccccttcatcagcgccggcgactatgaggggctgg
aaatgaaatccacccggtatctccagacccagggcagtgtgtacaaagtcagccgcctggagaccaccatctccacagtggtgggggcccac
gaggaggagccagaggacggccccaaggccacaccctcgtccctggacctgacctccaactgctcttcacgaagtgactccaagaccatga

cagagagcttcagcttctcctccaatgtgctctccgattacaaggatgacgacgataagtgataaacccgctgatcagcctcgactgtgccttctag
ttgccagccatctgttgtttgcccctcccccgtgccttccttgaccctggaaggtgccactcccactgtcctttcctaataaaatgaggaaattgcatcg
cattgtctgagtaggtgtcattctattctggggggtggggtggggcaggacagcaagggggaggattgggaagacaatagcaggcatgctggg
gatgcggtgggctctatggcttctgaggcggaaagaaccagctggggctctagggggtatccccacgcgccctgtagcggcgcattaagcgcg
gcgggtgtggtggttacgcgcagcgtgaccgctacacttgccagcgccctagcgcccgctcctttcgctttcttcccttcctttctcgccacgttcgcc
ggctttccccgtcaagctctaaatcgggggctccctttagggttccgatttagtgctttacggcacctcgaccccaaaaaacttgattagggtgatggt
tcacgtagtgggccatcgccctgatagacggtttttcgccctttgacgttggagtccacgttctttaatagtggactcttgttccaaactggaacaacac
tcaaccctatctcggtctattcttttgatttataagggattttgccgatttcggcctattggttaaaaaatgagctgatttaacaaaaatttaacgcgaatta
attctgtggaatgtgtgtcagttagggtgtggaaagtccccaggctccccagcaggcagaagtatgcaaagcatgcatctcaattagtcagcaac
caggtgtggaaagtccccaggctccccagcaggcagaagtatgcaaagcatgcatctcaattagtcagcaaccatagtcccgcccctaactcc
gcccatcccgcccctaactccgcccagttccgcccattctccgccccatggctgactaattttttttatttatgcagaggccgaggccgcctctgcctct
gagctattccagaagtagtgaggaggcttttttggaggcctaggcttttgcaaaaagctcccgggagcttgtatatccattttcggatctgatcaaga
gacaggatgaggatcgtttcgcatgattgaacaagatggattgcacgcaggttctccggccgcttgggtggagaggctattcggctatgactggg
cacaacagacaatcggctgctctgatgccgccgtgttccggctgtcagcgcaggggcgcccggttctttttgtcaagaccgacctgtccggtgccc
tgaatgaactgcaggacgaggcagcgcggctatcgtggctggccacgacgggcgttccttgcgcagctgtgctcgacgttgtcactgaagcgg
gaagggactggctgctattgggcgaagtgccggggcaggatctcctgtcatctcaccttgctcctgccgagaaagtatccatcatggctgatgca
atgcggcggctgcatacgcttgatccggctacctgcccattcgaccaccaagcgaaacatcgcatcgagcgagcacgtactcggatggaagcc
ggtcttgtcgatcaggatgatctggacgaagagcatcaggggctcgcgccagccgaactgttcgccaggctcaaggcgcgcatgcccgacgg
cgaggatctcgtcgtgacccatggcgatgcctgcttgccgaatatcatggtggaaaatggccgcttttctggattcatcgactgtggccggctgggt
gtggcggaccgctatcaggacatagcgttggctacccgtgatattgctgaagagcttggcggcgaatgggctgaccgcttcctcgtgctttacggt
atcgccgctcccgattcgcagcgcatcgccttctatcgccttcttgacgagttcttctgagcgggactctggggttcgaaatgaccgaccaagcgac
gcccaacctgccatcacgagatttcgattccaccgccgccttctatgaaaggttgggcttcggaatcgttttccgggacgccggctggatgatcctc
cagcgcggggatctcatgctggagttcttcgcccaccccaacttgtttattgcagcttataatggttacaaataaagcaatagcatcacaaatttcac
aaataaagcatttttttcactgcattctagttgtggtttgtccaaactcatcaatgtatcttatcatgtctgtataccgtcgacctctagctagagcttggcgt
aatcatggtcatagctgtttcctgtgtgaaattgttatccgctcacaattccacacaacatacgagccggaagcataaagtgtaaagcctggggtgc
ctaatgagtgagctaactcacattaattgcgttgcgctcactgcccgctttccagtcgggaaacctgtcgtgccagctgcattaatgaatcggccaa
cgcgcggggagaggcggtttgcgtattgggcgctcttccgcttcctcgctcactgactcgctgcgctcggtcgttcggctgcggcgagcggtatca
gctcactcaaaggcggtaatacggttatccacagaatcaggggataacgcaggaaagaacatgtgagcaaaaggccagcaaaaggccag
gaaccgtaaaaaggccgcgttgctggcgtttttccataggctccgcccccctgacgagcatcacaaaaatcgacgctcaagtcagaggtggcg
aaacccgacaggactataaagataccaggcgtttccccctggaagctccctcgtgcgctctcctgttccgaccctgccgcttaccggatacctgtc
cgcctttctcccttcgggaagcgtggcgctttctcatagctcacgctgtaggtatctcagttcggtgtaggtcgttcgctccaagctgggctgtgtgcac
gaaccccccgttcagcccgaccgctgcgccttatccggtaactatcgtcttgagtccaacccggtaagacacgacttatcgccactggcagcagc
cactggtaacaggattagcagagcgaggtatgtaggcggtgctacagagttcttgaagtggtggcctaactacggctacactagaagaacagta
tttggtatctgcgctctgctgaagccagttaccttcggaaaaagagttggtagctcttgatccggcaaacaaaccaccgctggtagcggtggtttttt
gtttgcaagcagcagattacgcgcagaaaaaaaggatctcaagaagatcctttgatcttttctacggggtctgacgctcagtggaacgaaaactc
acgttaagggattttggtcatgagattatcaaaaaggatcttcacctagatccttttaaattaaaaatgaagttttaaatcaatctaaagtatatatgagt
aaacttggtctgacagttaccaatgcttaatcagtgaggcacctatctcagcgatctgtctatttcgttcatccatagttgcctgactccccgtcgtgtag
ataactacgatacgggagggcttaccatctggccccagtgctgcaatgataccgcgagacccacgctcaccggctccagatttatcagcaataa
accagccagccggaagggccgagcgcagaagtggtcctgcaactttatccgcctccatccagtctattaattgttgccgggaagctagagtaagt
agttcgccagttaatagtttgcgcaacgttgttgccattgctacaggcatcgtggtgtcacgctcgtcgtttggtatggcttcattcagctccggttccca
acgatcaaggcgagttacatgatcccccatgttgtgcaaaaaagcggttagctccttcggtcctccgatcgttgtcagaagtaagttggccgcagt
gttatcactcatggttatggcagcactgcataattctcttactgtcatgccatccgtaagatgcttttctgtgactggtgagtactcaaccaagtcattctg
agaatagtgtatgcggcgaccgagttgctcttgcccggcgtcaatacgggataataccgcgccacatagcagaactttaaaagtgctcatcattg
gaaaacgttcttcggggcgaaaactctcaaggatcttaccgctgttgagatccagttcgatgtaacccactcgtgcacccaactgatcttcagcatc
ttttactttcaccagcgtttctgggtgagcaaaaacaggaaggcaaaatgccgcaaaaaagggaataagggcgacacggaaatgttgaatact
catactcttcctttttcaatattattgaagcatttatcagggttattgtctcatgagcggatacatatttgaatgtatttagaaaaataaacaaataggggtt
ccgcgcacatttccccgaaaagtgccacctgacgtc

Wild-type NK1R (receptor sequence underlined)

gacggatcgggagatctcccgatcccctatggtgcactctcagtacaatctgctctgatgccgcatagttaagccagtatctgctccctgcttgtgtgt
tggaggtcgctgagtagtgcgcgagcaaaatttaagctacaacaaggcaaggcttgaccgacaattgcatgaagaatctgcttagggttaggcg
ttttgcgctgcttcgcgatgtacgggccagatatacgcgttgacattgattattgactagttattaatagtaatcaattacggggtcattagttcatagcc
catatatggagttccgcgttacataacttacggtaaatggcccgcctggctgaccgcccaacgacccccgcccattgacgtcaataatgacgtat
gttcccatagtaacgccaatagggactttccattgacgtcaatgggtggagtatttacggtaaactgcccacttggcagtacatcaagtgtatcatat
gccaagtacgccccctattgacgtcaatgacggtaaatggcccgcctggcattatgcccagtacatgaccttatgggactttcctacttggcagtac
atctacgtattagtcatcgctattaccatggtgatgcggttttggcagtacatcaatgggcgtggatagcggtttgactcacggggatttccaagtctcc
accccattgacgtcaatgggagtttgttttggcaccaaaatcaacgggactttccaaaatgtcgtaacaactccgccccattgacgcaaatgggcg
gtaggcgtgtacggtgggaggtctatataagcagagctctctggctaactagagaacccactgcttactggcttatcgaaattaatacgactcacta
tagggagacccaagctggctagcgtttaaacttaagcttggtaccgagctcggatccgccaccatggataacgtcctcccggtggactcagacct
ctccccaaacatctccactaacacctcggaacccaatcagttcgtgcaaccagcctggcaaattgtcctttgggcagctgcctacacggtcattgt
ggtgacctctgtggtgggcaacgtggtagtgatgtggatcatcttagcccacaaaagaatgaggacagtgacgaactattttctggtgaacctggc
cttcgcggaggcctccatggctgcattcaatacagtggtgaacttcacctatgctgtccacaacgaatggtactacggcctgttctactgcaagttcc
acaacttctttcccatcgccgctgtcttcgccagtatctactccatgacggctgtggcctttgataggtacatggccatcatacatcccctccagcccc
ggctgtcagccacagccaccaaagtggtcatctgtgtcatctgggtcctggctctcctgctggccttcccccagggctactactcaaccacagaga
ccatgcccagcagagtcgtgtgcatgatcgaatggccagagcatccgaacaagatttatgagaaagtgtaccacatctgtgtgactgtgctgatct
acttcctcccccctgctggtgattggctatgcatacaccgtagtgggaatcacactatgggccagtgagatccccggggactcctctgaccgctacc
acgagcaagtctctgccaagcgcaaggtggtcaaaatgatgattgtcgtggtgtgcaccttcgccatctgctggctgcccttccacatcttcttcctcc
tgccctacatcaacccagatctctacctgaagaagtttatccagcaggtctacctggccatcatgtggctggccatgagctccaccatgtacaacc
ccatcatctactgctgcctcaatgacaggttccgtctgggcttcaagcatgccttccggtgctgccccttcatcagcgccggcgactatgaggggct
ggaaatgaaatccacccggtatctccagacccagggcagtgtgtacaaagtcagccgcctggagaccaccatctccacagtggtgggggccc
acgaggaggagccagaggacggccccaaggccacaccctcgtccctggacctgacctccaactgctcttcacgaagtgactccaagaccat
gacagagagcttcagcttctcctccaatgtgctctccgattacaaggatgacgacgataagtgataaacccgctgatcagcctcgactgtgccttct
agttgccagccatctgttgtttgcccctcccccgtgccttccttgaccctggaaggtgccactcccactgtcctttcctaataaaatgaggaaattgcat
cgcattgtctgagtaggtgtcattctattctggggggtggggtggggcaggacagcaagggggaggattgggaagacaatagcaggcatgctg
gggatgcggtgggctctatggcttctgaggcggaaagaaccagctggggctctagggggtatccccacgcgccctgtagcggcgcattaagcg
cggcgggtgtggtggttacgcgcagcgtgaccgctacacttgccagcgccctagcgcccgctcctttcgctttcttcccttcctttctcgccacgttcg
ccggctttccccgtcaagctctaaatcgggggctccctttagggttccgatttagtgctttacggcacctcgaccccaaaaaacttgattagggtgat
ggttcacgtagtgggccatcgccctgatagacggtttttcgccctttgacgttggagtccacgttctttaatagtggactcttgttccaaactggaacaa
cactcaaccctatctcggtctattcttttgatttataagggattttgccgatttcggcctattggttaaaaaatgagctgatttaacaaaaatttaacgcga
attaattctgtggaatgtgtgtcagttagggtgtggaaagtccccaggctccccagcaggcagaagtatgcaaagcatgcatctcaattagtcagc
aaccaggtgtggaaagtccccaggctccccagcaggcagaagtatgcaaagcatgcatctcaattagtcagcaaccatagtcccgcccctaa
ctccgcccatcccgcccctaactccgcccagttccgcccattctccgccccatggctgactaattttttttatttatgcagaggccgaggccgcctctg
cctctgagctattccagaagtagtgaggaggcttttttggaggcctaggcttttgcaaaaagctcccgggagcttgtatatccattttcggatctgatca
agagacaggatgaggatcgtttcgcatgattgaacaagatggattgcacgcaggttctccggccgcttgggtggagaggctattcggctatgact
gggcacaacagacaatcggctgctctgatgccgccgtgttccggctgtcagcgcaggggcgcccggttctttttgtcaagaccgacctgtccggt
gccctgaatgaactgcaggacgaggcagcgcggctatcgtggctggccacgacgggcgttccttgcgcagctgtgctcgacgttgtcactgaag
cgggaagggactggctgctattgggcgaagtgccggggcaggatctcctgtcatctcaccttgctcctgccgagaaagtatccatcatggctgat
gcaatgcggcggctgcatacgcttgatccggctacctgcccattcgaccaccaagcgaaacatcgcatcgagcgagcacgtactcggatgga
agccggtcttgtcgatcaggatgatctggacgaagagcatcaggggctcgcgccagccgaactgttcgccaggctcaaggcgcgcatgcccg
acggcgaggatctcgtcgtgacccatggcgatgcctgcttgccgaatatcatggtggaaaatggccgcttttctggattcatcgactgtggccggct
gggtgtggcggaccgctatcaggacatagcgttggctacccgtgatattgctgaagagcttggcggcgaatgggctgaccgcttcctcgtgcttta
cggtatcgccgctcccgattcgcagcgcatcgccttctatcgccttcttgacgagttcttctgagcgggactctggggttcgaaatgaccgaccaag
cgacgcccaacctgccatcacgagatttcgattccaccgccgccttctatgaaaggttgggcttcggaatcgttttccgggacgccggctggatga
tcctccagcgcggggatctcatgctggagttcttcgcccaccccaacttgtttattgcagcttataatggttacaaataaagcaatagcatcacaaatt
tcacaaataaagcatttttttcactgcattctagttgtggtttgtccaaactcatcaatgtatcttatcatgtctgtataccgtcgacctctagctagagcttg
gcgtaatcatggtcatagctgtttcctgtgtgaaattgttatccgctcacaattccacacaacatacgagccggaagcataaagtgtaaagcctggg
gtgcctaatgagtgagctaactcacattaattgcgttgcgctcactgcccgctttccagtcgggaaacctgtcgtgccagctgcattaatgaatcggc
caacgcgcggggagaggcggtttgcgtattgggcgctcttccgcttcctcgctcactgactcgctgcgctcggtcgttcggctgcggcgagcggta
tcagctcactcaaaggcggtaatacggttatccacagaatcaggggataacgcaggaaagaacatgtgagcaaaaggccagcaaaaggcc
aggaaccgtaaaaaggccgcgttgctggcgtttttccataggctccgcccccctgacgagcatcacaaaaatcgacgctcaagtcagaggtgg

cgaaacccgacaggactataaagataccaggcgtttccccctggaagctccctcgtgcgctctcctgttccgaccctgccgcttaccggatacctg
tccgcctttctcccttcgggaagcgtggcgctttctcatagctcacgctgtaggtatctcagttcggtgtaggtcgttcgctccaagctgggctgtgtgc
acgaaccccccgttcagcccgaccgctgcgccttatccggtaactatcgtcttgagtccaacccggtaagacacgacttatcgccactggcagca
gccactggtaacaggattagcagagcgaggtatgtaggcggtgctacagagttcttgaagtggtggcctaactacggctacactagaagaaca
gtatttggtatctgcgctctgctgaagccagttaccttcggaaaaagagttggtagctcttgatccggcaaacaaaccaccgctggtagcggtggttt
ttttgtttgcaagcagcagattacgcgcagaaaaaaaggatctcaagaagatcctttgatcttttctacggggtctgacgctcagtggaacgaaaa
ctcacgttaagggattttggtcatgagattatcaaaaaggatcttcacctagatccttttaaattaaaaatgaagttttaaatcaatctaaagtatatatg
agtaaacttggtctgacagttaccaatgcttaatcagtgaggcacctatctcagcgatctgtctatttcgttcatccatagttgcctgactccccgtcgtg
tagataactacgatacgggagggcttaccatctggccccagtgctgcaatgataccgcgagacccacgctcaccggctccagatttatcagcaa
taaaccagccagccggaagggccgagcgcagaagtggtcctgcaactttatccgcctccatccagtctattaattgttgccgggaagctagagta
agtagttcgccagttaatagtttgcgcaacgttgttgccattgctacaggcatcgtggtgtcacgctcgtcgtttggtatggcttcattcagctccggttc
ccaacgatcaaggcgagttacatgatcccccatgttgtgcaaaaaagcggttagctccttcggtcctccgatcgttgtcagaagtaagttggccgc
agtgttatcactcatggttatggcagcactgcataattctcttactgtcatgccatccgtaagatgcttttctgtgactggtgagtactcaaccaagtcatt
ctgagaatagtgtatgcggcgaccgagttgctcttgcccggcgtcaatacgggataataccgcgccacatagcagaactttaaaagtgctcatca
ttggaaaacgttcttcggggcgaaaactctcaaggatcttaccgctgttgagatccagttcgatgtaacccactcgtgcacccaactgatcttcagc
atcttttactttcaccagcgtttctgggtgagcaaaaacaggaaggcaaaatgccgcaaaaaagggaataagggcgacacggaaatgttgaat
actcatactcttcctttttcaatattattgaagcatttatcagggttattgtctcatgagcggatacatatttgaatgtatttagaaaaataaacaaatagg
ggttccgcgcacatttccccgaaaagtgccacctgacgtc

$\underline{\text{Nb}_{6e}}$

tggcgaatgggacgcgccctgtagcggcgcattaagcgcggcgggtgtggtggttacgcgcagcgtgaccgctacacttgccagcg
ccctagcgcccgctcctttcgctttcttcccttcctttctcgccacgttcgccggctttccccgtcaagctctaaatcgggggctccctttagg
gttccgatttagtgctttacggcacctcgaccccaaaaaacttgattagggtgatggttcacgtagtgggccatcgccctgatagacggtt
tttcgccctttgacgttggagtccacgttctttaatagtggactcttgttccaaactggaacaacactcaaccctatctcggtctattcttttgatt
tataagggattttgccgatttcggcctattggttaaaaaatgagctgatttaacaaaaatttaacgcgaattttaacaaaatattaacgttta
caatttcaggtggcacttttcggggaaatgtgcgcggaacccctatttgtttatttttctaaatacattcaaatatgtatccgctcatgaattaa
ttcttagaaaaactcatcgagcatcaaatgaaactgcaatttattcatatcaggattatcaataccatatttttgaaaaagccgtttctgtaat
gaaggagaaaactcaccgaggcagttccataggatggcaagatcctggtatcggtctgcgattccgactcgtccaacatcaatacaa
cctattaatttcccctcgtcaaaaataaggttatcaagtgagaaatcaccatgagtgacgactgaatccggtgagaatggcaaaagttt
atgcatttctttccagacttgttcaacaggccagccattacgctcgtcatcaaaatcactcgcatcaaccaaaccgttattcattcgtgattg
cgcctgagcgagacgaaatacgcgatcgctgttaaaaggacaattacaaacaggaatcgaatgcaaccggcgcaggaacactgc
cagcgcatcaacaatattttcacctgaatcaggatattcttctaatacctggaatgctgttttcccggggatcgcagtggtgagtaaccatg
catcatcaggagtacggataaaatgcttgatggtcggaagaggcataaattccgtcagccagtttagtctgaccatctcatctgtaacat
cattggcaacgctacctttgccatgtttcagaaacaactctggcgcatcgggcttcccatacaatcgatagattgtcgcacctgattgccc
gacattatcgcgagcccatttatacccatataaatcagcatccatgttggaatttaatcgcggcctagagcaagacgtttcccgttgaata
tggctcataacaccccttgtattactgtttatgtaagcagacagttttattgttcatgaccaaaatcccttaacgtgagttttcgttccactgag
cgtcagaccccgtagaaaagatcaaaggatcttcttgagatcctttttttctgcgcgtaatctgctgcttgcaaacaaaaaaaccaccgc
taccagcggtggtttgtttgccggatcaagagctaccaactctttttccgaaggtaactggcttcagcagagcgcagataccaaatactg
tccttctagtgtagccgtagttaggccaccacttcaagaactctgtagcaccgcctacatacctcgctctgctaatcctgttaccagtggct
gctgccagtggcgataagtcgtgtcttaccgggttggactcaagacgatagttaccggataaggcgcagcggtcgggctgaacggg
gggttcgtgcacacagcccagcttggagcgaacgacctacaccgaactgagatacctacagcgtgagctatgagaaagcgccacg
cttcccgaagggagaaaggcggacaggtatccggtaagcggcagggtcggaacaggagagcgcacgagggagcttccagggg
gaaacgcctggtatctttatagtcctgtcgggtttcgccacctctgacttgagcgtcgatttttgtgatgctcgtcaggggggcggagcctat
ggaaaaacgccagcaacgcggcctttttacggttcctggccttttgctggccttttgctcacatgttctttcctgcgttatcccctgattctgtg
gataaccgtattaccgcctttgagtgagctgataccgctcgccgcagccgaacgaccgagcgcagcgagtcagtgagcgaggaag
cggaagagcgcctgatgcggtattttctccttacgcatctgtgcggtatttcacaccgcatatatggtgcactctcagtacaatctgctctga

tgccgcatagttaagccagtatacactccgctatcgctacgtgactgggtcatggctgcgccccgacacccgccaacacccgctgac
gcgccctgacgggcttgtctgctcccggcatccgcttacagacaagctgtgaccgtctccgggagctgcatgtgtcagaggttttcacc
gtcatcaccgaaacgcgcgaggcagctgcggtaaagctcatcagcgtggtcgtgaagcgattcacagatgtctgcctgttcatccgcg
tccagctcgttgagtttctccagaagcgttaatgtctggcttctgataaagcgggccatgttaagggcggttttttcctgtttggtcactgatgc
ctccgtgtaagggggatttctgttcatgggggtaatgataccgatgaaacgagagaggatgctcacgatacgggttactgatgatgaa
catgcccggttactggaacgttgtgagggtaaacaactggcggtatggatgcggcgggaccagagaaaaatcactcagggtcaatg
ccagcgcttcgttaatacagatgtaggtgttccacagggtagccagcagcatcctgcgatgcagatccggaacataatggtgcaggg
cgctgacttccgcgtttccagactttacgaaacacggaaaccgaagaccattcatgttgttgctcaggtcgcagacgttttgcagcagca
gtcgcttcacgttcgctcgcgtatcggtgattcattctgctaaccagtaaggcaaccccgccagcctagccgggtcctcaacgacagga
gcacgatcatgcgcacccgtggggccgccatgccggcgataatggcctgcttctcgccgaaacgtttggtggcgggaccagtgacg
aaggcttgagcgagggcgtgcaagattccgaataccgcaagcgacaggccgatcatcgtcgcgctccagcgaaagcggtcctcgc
cgaaaatgacccagagcgctgccggcacctgtcctacgagttgcatgataaagaagacagtcataagtgcggcgacgatagtcatg
ccccgcgcccaccggaaggagctgactgggttgaaggctctcaagggcatcggtcgagatcccggtgcctaatgagtgagctaactt
acattaattgcgttgcgctcactgcccgctttccagtcgggaaacctgtcgtgccagctgcattaatgaatcggccaacgcgcggggag
aggcggtttgcgtattgggcgccagggtggtttttcttttcaccagtgagacgggcaacagctgattgcccttcaccgcctggccctgag
agagttgcagcaagcggtccacgctggtttgccccagcaggcgaaaatcctgtttgatggtggttaacggcgggatataacatgagct
gtcttcggtatcgtcgtatcccactaccgagatatccgcaccaacgcgcagcccggactcggtaatggcgcgcattgcgcccagcgc
catctgatcgttggcaaccagcatcgcagtgggaacgatgccctcattcagcatttgcatggtttgttgaaaaccggacatggcactcc
agtcgccttcccgttccgctatcggctgaatttgattgcgagtgagatatttatgccagccagccagacgcagacgcgccgagacaga
acttaatgggcccgctaacagcgcgatttgctggtgacccaatgcgaccagatgctccacgcccagtcgcgtaccgtcttcatgggag
aaaataatactgttgatgggtgtctggtcagagacatcaagaaataacgccggaacattagtgcaggcagcttccacagcaatggca
tcctggtcatccagcggatagttaatgatcagcccactgacgcgttgcgcgagaagattgtgcaccgccgctttacaggcttcgacgcc
gcttcgttctaccatcgacaccaccacgctggcacccagttgatcggcgcgagatttaatcgccgcgacaatttgcgacggcgcgtgc
agggccagactggaggtggcaacgccaatcagcaacgactgtttgcccgccagttgttgtgccacgcggttgggaatgtaattcagct
ccgccatcgccgcttccactttttcccgcgttttcgcagaaacgtggctggcctggttcaccacgcgggaaacggtctgataagagaca
ccggcatactctgcgacatcgtataacgttactggtttcacattcaccaccctgaattgactctcttccgggcgctatcatgccataccgcg
aaaggttttgcgccattcgatggtgtccgggatctcgacgctctcccttatgcgactcctgcattaggaagcagcccagtagtaggttga
ggccgttgagcaccgccgccgcaaggaatggtgcatgcaaggagatggcgcccaacagtcccccggccacggggcctgccacc
atacccacgccgaaacaagcgctcatgagcccgaagtggcgagcccgatcttccccatcggtgatgtcggcgatataggcgccag
caaccgcacctgtggcgccggtgatgccggccacgatgcgtccggcgtagaggatcgagatctcgatcccgcgaaattaatacgac
tcactataggggaattgtgagcggataacaattcccctctagaaataattttgtttaactttaagaaggagatatacatatgaaatacctg
ctgccgaccgctgctgctggtctgctgctcctcgctgcccagccggcgatggcccaagtccaattacaagagtccggcggcggacttg
tccagcctggggggatcattgcgcctgtcgtgtgcggcgtcgggatttgtatttgaaaatagtgccatggcctggtatcgtcaggcgcctgg
gcgtgagcgtgagctgattgctgtgattggaactacattcattcgtttggctgaatctgtgcgtgggcgttttacgatcagccgcgataatg
cacgttctacggtttatttgcaaatgaataatttacgtcctgaagacacagcggtttactactgttcaaaatcgggagcgtattggggaca
ggggacacaagtaactgtgtcatctggcgggctgcccgagacgggcggccaccaccaccaccaccactgagatccggctgctaac
aaagcccgaaaggaagctgagttggctgctgccaccgctgagcaataactagcataaccccttggggcctctaaacgggtcttgag
gggttttttgctgaaaggaggaactatatccggat

$\underline{\text{Nb}_{\text{alfa}}}$

nagcgcccaatacgcaaaccgcctctccccgcgcgttggccgattcattaatgcagctggcacgacaggtttcccgactggaaagcg
ggcagtgagcgcaacgcaattaatgtgagttagctcactcattaggcaccccaggctttacactttatgcttccggctcgtatgttgtgtgg
aattgtgagcggataacaatttcacacaggaaacagctatgaccatgattacgccaagcttgcatgcaaattctatttcaaggagaca

gtcataatgaaatacctattgcctacggcagccgctggattgttattactcgcggcccagccggccatggctgaagtacagcttcagga
atccggtggcggtctggtacagccgggtggctccctgcgtctgagttgtaccgctagcggtgttaccatttctgccttaaacgccatggcg
atgggctggtatcgccaagcgccgggagaacgtcgcgtcatggtagcggccgtatcggagcgtggcaatgcgatgtaccgtgaaag
cgtacaagggcgctttacggtaacgcgcgacttcacgaacaaaatggtaagcttacaaatggataatctgaagccggaagacacg
gcagtgtattactgtcacgtattggaggatcgcgttgactctttccatgattattgggggcaaggtacacaggtgacggtatcctcagtctc
ctcaggaggactgccggaaaccggcggccaccaccatcaccatcactaatagaattcactggccgtcgttttacaacgtcgtgactg
ggaaaaccctggcgttacccaacttaatcgccttgcagcacatccccctttcgccagctggcgtaatagcgaagaggcccgcaccga
tcgcccttcccaacagttgcgcagcctgaatggcgaatggcgcctgatgcggtattttctccttacgcatctgtgcggtatttcacaccgc
atacgtcaaagcaaccatagtacgcgccctgtagcggcgcattaagcgcggcgggtgtggtggttacgcgcagcgtgaccgctaca
cttgccagcgccctagcgcccgctcctttcgctttcttcccttcctttctcgccacgttcgccggctttccccgtcaagctctaaatcgggggc
tccctttagggttccgatttagtgctttacggcacctcgaccccaaaaaacttgatttgggtgatggttcacgtagtgggccatcgccctga
tagacggtttttcgccctttgacgttggagtccacgttctttaatagtggactcttgttccaaactggaacaacactcaaccctatctcgggct
attcttttgatttataagggattttgccgatttcggcctattggttaaaaaatgagctgatttaacaaaaatttaacgcgaattttaacaaaata
ttaacgtttacaattttatggtgcactctcagtacaatctgctctgatgccgcatagttaagccagccccgacacccgccaacacccgctg
acgcgccctgacgggcttgtctgctcccggcatccgcttacagacaagctgtgaccgtctccgggagctgcatgtgtcagaggttttca
ccgtcatcaccgaaacgcgcgagacgaaagggcctcgtgatacgcctatttttataggttaatgtcatgataataatggtttcttagacgt
caggtggcacttttcggggaaatgtgcgcggaacccctatttgtttatttttctaaatacattcaaatatgtatccgctcatgagacaataac
cctgataaatgcttcaataatattgaaaaaggaagagtatgagtattcaacatttccgtgtcgcccttattcccttttttgcggcattttgccttc
ctgtttttgctcacccagaaacgctggtgaaagtaaaagatgctgaagatcagttgggtgcacgagtgggttacatcgaactggatctc
aacagcggtaagatccttgagagttttcgccccgaagaacgttttccaatgatgagcacttttaaagttctgctatgtggcgcggtattatc
ccgtattgacgccgggcaagagcaactcggtcgccgcatacactattctcagaatgacttggttgagtactcaccagtcacagaaaa
gcatcttacggatggcatgacagtaagagaattatgcagtgctgccataaccatgagtgataacactgcggccaacttacttctgacaa
cgatcggaggaccgaaggagctaaccgcttttttgcacaacatgggggatcatgtaactcgccttgatcgttgggaaccggagctgaa
tgaagccataccaaacgacgagcgtgacaccacgatgcctgtagcaatggcaacaacgttgcgcaaactattaactggcgaacta
cttactctagcttcccggcaacaattaatagactggatggaggcggataaagttgcaggaccacttctgcgctcggcccttccggctgg
ctggtttattgctgataaatctggagccggtgagcgtgggtctcgcggtatcattgcagcactggggccagatggtaagccctcccgtat
cgtagttatctacacgacggggagtcaggcaactatggatgaacgaaatagacagatcgctgagataggtgcctcactgattaagca
ttggtaactgtcagaccaagtttactcatatatactttagattgatttaaaacttcatttttaatttaaaaggatctaggtgaagatcctttttgat
aatctcatgaccaaaatcccttaacgtgagttttcgttccactgagcgtcagaccccgtagaaaagatcaaaggatcttcttgagatcctt
tttttctgcgcgtaatctgctgcttgcaaacaaaaaaaccaccgctaccagcggtggtttgtttgccggatcaagagctaccaactctttttc
cgaaggtaactggcttcagcagagcgcagataccaaatactgtccttctagtgtagccgtagttaggccaccacttcaagaactctgta
gcaccgcctacatacctcgctctgctaatcctgttaccagtggctgctgccagtggcgataagtcgtgtcttaccgggttggactcaaga
cgatagttaccggataaggcgcagcggtcgggctgaacggggggttcgtgcacacagcccagcttggagcgaacgacctacacc
gaactgagatacctacagcgtgagctatgagaaagcgccacgcttcccgaagggagaaaggcggacaggtatccggtaagcggc
agggtcggaacaggagagcgcacgagggagcttccagggggaaacgcctggtatctttatagtcctgtcgggtttcgccacctctga
cttgagcgtcgatttttgtgatgctcgtcaggggggcggagcctatggaaaaacgccagcaacgcggcctttttacggttcctggcctttt
gctggccttttgctcacatgttctttcctgcgttatcccctgattctgtggataaccgtattaccgcctttgagtgagctgataccgctcgccgc
agccgaacgaccgagcgcagcgagtcagtgagcgaggaagcggaag

NK1R sequence

GPSRLEEELRRRLTEPGQADQEAKELARQISGPDRVRAVSHWSSMDNVLPVDSDLSPNISTNT
SEPNQFVQPAWQIVLWAAAYTVIVVTSVVGNVVVMWIILAHKRMRTVTNYFLVNLAFAEASMAA
FNTVVNFTYAVHNEWYYGLFYCKFHNFFPIAAVFASIYSMTAVAFDRYMAIIHPLQPRLSATATK

VVICVIWVLALLLAFPQGYYSTTETMPSRVVCMIEWPEHPNKIYEKVYHICVTVLIYFLPLLVIGYA
YTVVGITLWASEIPGDSSDRYHEQVSAKRKVVKMMIVVVCTFAICWLPFHIFFLLPYINPDLYLKK
FIQQVYLAIMWLAMSSTMYNPIIYCCLNDRFRLGFKHAFRCCPFISAGDYEGLEMKSTRYLQTQ
GSVYKVSRLETTISTVVGAHEEEPEDGPKATPSSLDLTSNCSSRSDSKTMTESFSFSSNVLSDY
KDDDDK

Highlighted alfa, 6e and $BC_2$ tags

Nb sequences

$Nb_{6e}$: QVQLQESGGG LVQPGGSLRL SCAASGFVFE NSAMAWYRQA PGKERELIAV IGTTFIKLAE SVKGRFTISR DNAKSTVYLQ MNNLKPEDTA VYYCSKSGAY WGQGTQVTVS SGGLPETGHH HHHH

$Nb_{alfa}$: EVQLQESGGG LVQPGGSLRL SCTASGVTIS ALNAMAMGWY RQAPGERRVM VAAVSERGNA MYRESVQGRF TVTRDFTNKM VSLQMDNLKP EDTAVYYCHV LEDRVDSFHD YWGQGTQVTV SSGGLPETGG HHHHHH

$Nb_{BC2}$: QVQLVESGGG LVQPGGSLTL SCTASGFTLD HYDIGWFRQA PGKEREGVSC INNSDDDTYY ADSVKGRFTI FMNNAKDTVY LQMNSLKPED TAIYYCAEAR GCKRGRYEYD FWGQGTQVTV SSGGLPETGG HHHHHH

$Nb_{GFP}$: QVQLQESGGA LVQPGGSLRL SCAASGFPVN RYSMRWYRQA PGKEREWVAG MSSAGDRSSY EDSVKGRFTI SRDDARNTVY LQMNSLKPED TAVYYCNVNV GFEYWGQGTQ VTVSSGGLPE TGGHHHHHH

**Table S1.** **Nanobody-epitope tag pairs**

| Tag | Sequence |
|---|---|
| 6e | QADQEAKELARQIS |
| Alfa | SRLEEELRRRLTE |
| $BC_2$ | PDRVRAVSHWSS |

**Table S2. List of Nb conjugates calculated and observed masses**

| Nb conjugates | calculated mass after sortagging | observed mass |
|---|---|---|
| $Nb_{alfa}$-LPTGG-$His_6$ | - | 15207 |
| $Nb_{6e}$-LPTGG-$His_6$ | - | 13476 |
| $Nb_{BC2}$-LPTGG-$His_6$ | - | 14993 |
| $Nb_{GFP}$-LPTGG-$His_6$ | - | 14233 |
| $Nb_{alfa}$-NKA | 15556 | 15555 |
| $Nb_{6e}$-NKA | 13825 | 13828 |
| $Nb_{BC2}$-NKA | 15342 | 15338 |
| $Nb_{GFP}$-NKA | 14582 | 14583 |
| $Nb_{alfa}$-$SP_{6-11}$ | 15164 | 15165 |
| $Nb_{6e}$-$SP_{6-11}$ | 13433 | 13436 |
| $Nb_{BC2}$-$SP_{6-11}$ | 14950 | 14951 |
| $Nb_{GFP}$-$SP_{6-11}$ | 14190 | 14189 |
